# Efficient CRISPR/Cas9-based genome editing and its application to conditional genetic analysis in *Marchantia polymorpha*

**DOI:** 10.1101/277350

**Authors:** Shigeo S. Sugano, Ryuichi Nishihama, Makoto Shirakawa, Junpei Takagi, Yoriko Matsuda, Sakiko Ishida, Tomoo Shimada, Ikuko Hara-Nishimura, Keishi Osakabe, Takayuki Kohchi

## Abstract

*Marchantia polymorpha* is one of the model species of basal land plants. Although CRISPR/Cas9-based genome editing has already been demonstrated for this plant, the efficiency was too low to apply to functional analysis. In this study, we show the establishment of CRISPR/Cas9 genome editing vectors with high efficiency for both construction and genome editing. Codon optimization of Cas9 to Arabidopsis achieved over 70% genome editing efficiency at two loci tested. Systematic assessment revealed that guide sequences of 17 nt or shorter dramatically decreased this efficiency. We also demonstrated that a combinatorial use of this system and a floxed complementation construct enabled conditional analysis of a nearly essential gene. This study reports that simple, rapid, and efficient genome editing is feasible with the series of developed vectors.

## Introduction

The clustered regularly interspaced short palindromic repeats (CRISPR)/ CRISPR-associated endonuclease 9 (Cas9)-based genome editing system is a groundbreaking technology in molecular genetics, which enables alterations of target sequences in the genome [1, 2]. The system consists of two components: a Cas9 protein, which has an RNA-guided endonuclease activity, and a single guide RNA (gRNA), which specifies a target sequence within the genome. The Cas9 protein from *Streptococcus pyogenes* binds to the DNA sequence “NGG,” which is known as the protospacer adjacent motif (PAM) sequence. The interaction between a Cas9 protein and a PAM sequence induces the interaction between a gRNA and its target DNA sequence. If a sufficient length of the gRNA matches with the target sequence, the nuclease domains of Cas9 become capable of cutting the phosphodiester bonds on both sides of the strands, which are located 3 bp upstream of the PAM sequence [3]. Once a double-strand break (DSB) occurs, the error-prone non-homologous end joining (NHEJ) repair pathway is activated and sometimes introduces indels or base substitutions randomly at the target site, which could result in disruption of the target locus with various alleles.

This simple CRISPR/Cas9 system has been reconstructed in a wide range of eukaryotes, and researchers are now able to use molecular genetics in the species that are suitable for the purposes of their specific biological research [4]. Previous studies demonstrated that the CRISPR/Cas9 system works in a variety of plant species, from algae to crops [5-7], which has greatly changed functional genetics in basic and applied plant research. Recent studies on the moss *Physcomitrella patens* [8, 9] and the green alga *Chlamydomonas reinhardtii* [10, 11] reported highly efficient genome editing methods using CRISPR-based genome editing. Compared to flowering plants,such haploid generation-dominant plant species are free from the transheterozygosity issues associated with diploidy or polyploidy [12-14], allowing isolation of pure mutant lines for analysis with relative ease. In the meanwhile, regardless of the ploidy, but especially for haploid species, genome editing techniques cannot be simply applied to essential genes as this leads to lethality; conditional approaches are required.

The liverwort *Marchantia polymorpha* is an emerging model species of land plants for studying plant evolution and gene function [15]. *M. polymorpha* has good features for the application of reverse genetics. Most vascular plants and mosses are known to have experienced two or more whole genome duplication events, which makes it difficult to analyze gene functions due to the presence of paralogous genes. Sequencing of the *M. polymorpha* genome revealed no sign of a whole genome duplication and accordingly there is low genetic redundancy in most regulatory genes, such as transcription factors and signaling components [16]. In addition, non-chimeric individuals can be easily obtained and propagated via gemmae that are derived from single cells by asexual reproduction in *M. polymorpha* [17], which accelerates transgenic experiments [18]. A variety of tools for molecular genetic experiments have been developed for *M. polymorpha* [18], such as high-efficiency transformation methods [19-21], a homologous recombination-mediated gene targeting method [22], a systematic set of vectors [18], and a conditional gene expression/deletion system [23].

Recently, a transcription activator-like effector nuclease (TALEN)-based genome editing technology was established in *M. polymorpha* [24]. We have previously demonstrated that a CRISPR/Cas9-based knockout system, which exploited human-codon-optimized Cas9, can operate in *M. polymorpha* [25]. However, the efficiency of genome editing was so low that the identification of plants with a mutation at the target locus required selection by a phenotype attributed to the mutation. Systematic optimization of Cas9 and gRNAs with rice, tobacco, and Arabidopsis showed that their expression levels greatly affected genome editing efficiencies [26, 27], suggesting that there is room to improve genome editing efficiencies in *M. polymorpha*.

Here, we report remarkable improvement in genome editing efficiency using Arabidopsis-codon-optimized (Atco) Cas9 to a degree that simple direct sequencing analysis of few number of transformants is sufficient to obtain genome-edited plants. In this efficient genome editing system, we assessed off-target effects and evaluated the influence of gRNAs length. Occurrence of large deletions using two gRNAs was also demonstrated. In addition, we provide a simple CRISPR/Cas9-based method to generate conditional knockout mutants. Our improved CRISPR/Cas9-based genome editing system can be used as a powerful molecular genetic tool in *M. polymorpha*.

## Materials and Methods

### Accessions, growth conditions and transformation of *M. polymorpha*

*M. polymorpha* Takaragaike-1 (Tak-1, male accession) and Takaragaike-2 (Tak-2, female accession) were used as wild types [19]. F1 spores were generated as previously described [25]. *M. polymorpha* was cultured axenically under 50–60 µmol m^−2^ sec^−1^ continuous white light at 22 °C. *Agrobacterium*-mediated transformation of F1 sporelings was performed as described previously [19]. Transformants were selected on half-strength B5 medium [28] containing 1% agar with 0.5 µM chlorsulfuron (kindly provided by DuPont; in case of the assay in Fig. 6, Wako Pure Chemical Industries) or 10 mg L^−1^ hygromycin (Wako Pure Chemical Industries) depending on the transformation vector.

### Vector construction

All the DNA sequences of the vectors were deposited to DDBJ and Addgene: pMpGE_En01(LC090754; 71534), pMpGE_En03 (LC090755; 71535), pMpGE010 (LC090756; 71536), pMpGE011 (LC090757; 71537), pMpGE006 (LC375817; 108722), pMpGE013 (LC375815; 108681), pMpGE014 (LC375816; 108682),pMpGWB337tdTN (LC375949; 108717), pMpGWB337Cit (LC375950; 108718),pMpGWB337tdT (LC375951; 108719), pMpGWB337TR (LC375952; 108720),pMpGWB337mT (LC375953; 108721). The construction of these vectors was performed as follows:

-pMpGE010, pMpGE011. Firstly, a DNA fragment of nuclear localization signal (NLS)-tagged *Atco-Cas9* with the *Pisum sativum rbcS3A* terminator (Pea3ter) was PCR amplified with the primers cacc_AtCas9_F and Pea3Ter_R using the pDe-CAS9 vector [12] as a template and subcloned into pENTR/D-TOPO (Invitrogen). Using the entry clone and LR Clonase II (Invitrogen), LR reactions with pMpGWB103 [29] and pMpGWB303 [29] were conducted to express Cas9 under the *M. polymorpha ELONGATION FACTOR1α* promoter (Mp*EF*_*pro*_) [18]. The LR reaction product using pMpGWB103 was designated pMpGE006 and used for the two vector system experiments (Fig. 1A). Next, a Gateway attR1-attR2 cassette amplified from pMpGWB303 using the primers Infusion_GW_A51_F and Infusion_GW_A51_R was subcloned into the AorHI51 restriction enzyme site of the vectors produced by the LR reaction above to produce pMpGE010 and pMpGE011.

**Fig 1.**
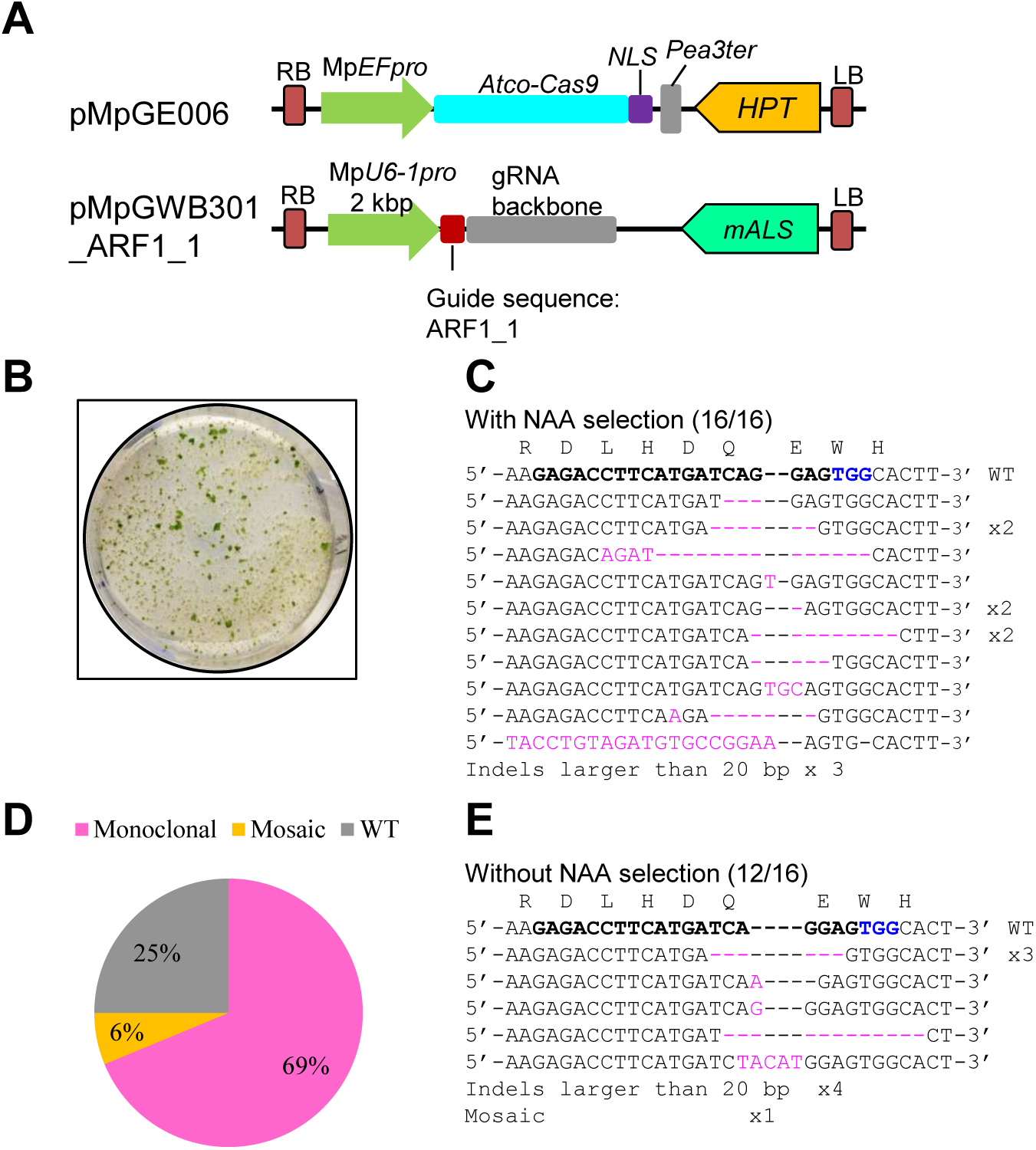
Improvement of the *M. polymorpha* CRISPR/Cas9 system by codon optimization. (A) Diagrams of the vectors used. pMpGE006 contains a cassette for the expression of Atco-Cas9 fused with an NLS under the control of Mp*EF*_*pro*_. pMpGWB301_ARF1_1 contains a cassette for the expression of the gRNA ARF1_1 under the control of Mp*U6-1_pro_* [25]. (B) Photograph of auxin-insensitive co-transformants of the two vectors described in (A). *Agrobacterium•*co-cultured sporelings that corresponded to one eighth of sporangium were plated on a medium in containing 3 µM NAA and two vector selection substances, 10 mg/L hygromycin and 0.5 µM chlorsulfuron. Diameter of the circle dish shows 10 cm. (C) Direct sequencing analysis of the target locus of Mp*ARF1* in auxin-selected T1 co-transformants. Inserted or substituted bases are colored in magenta. The target guide sequence of ARF1_1 is shown in bold face with the PAM sequence in blue. (D and E) Proportions of genome editing patterns (D) and sequences of the target site (E) in T1 co-transformants, which were obtained with no auxin selection. Sixteen independent transformants were analyzed for the target-site sequence by direct sequencing. “Monoclonal” and “Mosaic” indicate direct sequencing read patterns with mutated sequence peaks only and those with mixed sequence peaks, respectively. “WT” indicates read patterns identical to the original target sequence.

-pMpGE_En01. The 2 kbp promoter region of Mp*U6-1* was amplified from pENTR D-TOPO/MpU6-1pro:gRNA_ARF1 [25] using the primers Mp-U6_38003_F and Mp-U6_38003_R. The CmR-ccdB-gRNA fragment with SacI and PstI sites was also amplified from pENTR D-TOPO/AtU6pro:CmR-ccdB-gRNA (unpublished data) using the primers OE-MpU6-CmRccdB-F2 and gRNA-R3. These two amplified fragments were combined using overlap extension PCR and the combined fragment was cloned into pENTR/D-TOPO vector to produce pMpGE_En01.

-pMpGE_En02. PCR was conducted using pMpGE_En01 as a template and with the primers BsaI-Sp-sgRNA_F and gRNA_R to amplify a gRNA backbone fragment.Additionally, a 500 bp region of the Mp*U6-1* promoter was PCR-amplified using the MpU6-1_500_F and BsaI_MpU6_1R primers and plasmid pMpGE_En01 as a template. Amplified gRNA and Mp*U6-1* promoter fragments were conjugated by overlap extension PCR and cloned into pENTR/D-TOPO to produce pMpGE_En02.-MpGE_En03. pMpGE_En02 was digested by BsaI. Two oligo DNAs, Mp_oligo6BsaI_Gf and Mp_oligo6BsaI_Gr, were annealed and cloned into BsaI-digested pMpGE_En02 to produce the pMpGE_En03 plasmid (see Fig. S1).-pMpGE013 and pMpGE014 vectors. Briefly, annealed oligos harboring AarI recognition sites (Mp_oligo5AarI_Gf and Mp_oligo5AarI_Gr) were subjected to ligation reactions with the pMpGE_En02 vector linearized with BsaI. The resulting vector was subjected to LR reaction with pMpGE010 and pMpGE011 to produce pMpGE013 and pMpGE014, respectively.

-pMpGE vectors harboring *ARF1_1, NOP1_1*, and *NOP1_2* gRNA. pMpGWB301, harboring ARF1_1 gRNA, was the same construct as in the previous report [25].Construction of other vectors was by the methods described in Fig. S1. Briefly, annealed oligos of ARF1_1 and NOP1_1 for pMpGE_En01 were subjected to the In-Fusion^TM^ HD Cloning Kit (TaKaRa) to the pMpGE_En01 vector linearized with PstI and SacI (TaKaRa). Conversely, annealed oligos of NOP1_2 gRNAs were subjected to ligation reactions with pMpGE_En03 linearized with BsaI (NEB). These constructs were subjected to the LR reaction to be introduced into pMpGE010 or pMpGE011.Oligos for gRNAs are listed in Supplemental Table S1.

-pMpGWB337 series. The plasmid pMp301-EFp:loxGW:Tlox:CitNLS:T^18)^ was digested with SalI and SacI and ligated with a SalI-SacI fragment containing an NruI site, which was generated by PCR using pMp301-EFp:loxGW:Tlox:CitNLS:T as template with the primer set ccdB_236F and loxP_NruI_Sac_R and subsequent enzyme digestion, to generate pMp301-EFp:loxGW:Tlox-NruI. Coding sequences of various fluorescent proteins were PCR amplified using a common reverse primer (NOSt_head_R_SacI) and the following forward primers and templates: TagRFP (TagRFP_CAGC_F; pMpGWB126 [29]), tdTomato (tdTomato_CAGC_F; pMpGWB129 [29]), tdTomato-NLS (tdTomato_CAGC_F; pMpGWB116 [29]), Atco-mTurquoise2 (mTurq_CAGC_F, pUGW2-mTurq2 [see below]), and Citrine (ccdB_236F, pMpGWB337 [23]). These amplified fragments (except for Citrine) were digested with SacI and ligated with NruI/SacI-digested pMp301-EFp:loxGW:Tlox-NruI to generate a pMp301-EFp:loxGW:Tlox:FP:T series (FP: TR for TagRFP; tdT for tdTomato; tdTN for tdTomato-NLS; mT for mTurquoise2). For Citrine, the amplified fragment was digested with SalI and SacI and ligated with SalI/SacI-digested pMp301-EFp:loxGW:Tlox:CitNLS:T to generate pMp301-EFp:loxGW:Tlox:Cit:T. A fragment consisting of the Mp*HSP17.8A1* promoter, the Cre-GR coding sequence, and the NOS terminator was amplified and cloned into the AscI site of the pMp301-EFp:loxGW:Tlox:FP:T series, as described previously [29], to generate a series of pMpGWB337-FP vectors (Fig. S7). For construction of pUGW2-mTurq2, an Atco-mTurquoise2 DNA fragment, which was synthesized and amplified by PCR with the primer set pUGW_Aor_mTurq_IF_F and pUGW_Aor_mTurq_Stp_IF_R, was cloned into the unique Aor51HI site of pUGW2 [30] using the In-Fusion Cloning Kit. The primers used are listed in Supplementary Table S1.

### Mutation analyses in on-target and off-target sites

Transformed sporelings were selected with antibiotics for 18 days, and selected lines, referred to as T1 plants, were transferred to fresh medium with the same antibiotics for approximately 2 weeks. The genomic DNAs of T1 thalli were extracted. The Mp*ARF1* target locus was PCR amplified using the primers ARF1_Seq_F3 and ARF1_Seq_R3 and subjected to direct sequencing. The Mp*NOP1* target locus was PCR amplified using the primers CRISPR_NOP1_F and CRISPR_NOP1_R then subjected to direct sequencing. Using genomic DNAs harboring mutations in the on-target sites of ARF1_1 or NOP1_1, corresponding off-target sites were PCR amplified using the primers listed in Supplementary Table S1, and subjected to direct sequencing. Off-target sites of ARF1_1 and NOP1_1 were searched using the *M. polymorpha* genome ver3.1 [16] and CasOT software [31].

### Assessment of gRNA guide lengths

pMpGE010_NOP1_2 variants harboring various lengths of gRNAs were constructed using pMpGE_En02. For the addition of an “extra initial G,” the pMpGE_En03 vector was used (oligos are listed in Supplementary Table S1). T1 plants in petri dishes were placed onto a white-light-emitting display (iPad, Apple) and photos of the whole body of T1 plants were taken by a digital camera (EOS KissX3). Digital images of T1 plants were analysed by ImageJ. Using Threshold Colour program, ratio of transparent area to whole plant were measured. The images were classified into three types; Class I: over 90% of the thallus area was transparent; Class II: less than 90% and over 20% transparent area; and Class III: less than 20% transparent.

### Analysis of *de novo* mutations

Three pMpGE010_NOP1_1 transformants that had transparent (mutant) and non-transparent (wild-type) sectors in one individual were cultured. Four gemmae from a gemma cup formed on each of the two sectors were transplanted to new media and cultured to conduct genome analysis.

### Large deletion induction by co-transformation

pMpGE013 and pMpGE014 were digested by the AarI restriction enzyme. Annealed oligos for NOP1_3 to NOP1_6 (their sequences are described in Supplemental Table S1) were ligated into the linearized vectors. Vectors were introduced into regenerating thalli via *Agrobacterium* [20]. T1 transformants selected by appropriate antibiotics were cultured at for least 2 weeks and subjected to DNA extraction. The extracted DNAs from T1 thalli were analyzed by PCR using the appropriate primers described in Supplementary Table S1. The PCR products were analyzed by direct sequencing to examine genome-editing events.

### Generation of conditional knockout mutants

Annealed oligos for an Mp*MPK1*-targeting gRNA (sgRNA_Bsa_MpMPK1_ex1t_F and -R) were ligated into BsaI-digested pMpGE_En02. The resulting plasmid was subjected to LR reaction with pMpGE010 to generate pMpGE010-MPK1ex1t.Mp*MPK1* cDNA was amplified by RT-PCR using RNA from Tak-1 and the primer set MpMPK1_1F_TOPO and MpMPK1_c1131R_STP and cloned into pENTR/D-TOPO. The resulting entry vector was subjected to LR reaction with pMpGWB337 and pMpGWB337tdTN to generate pMpGWB337-cMPK1 and pMpGWB337tdTN-cMPK1, respectively [19]. Sporelings were transformed with pMpGE010-MPK1ex1t alone or together with pMpGWB337-cMPK1 or pMpGWB337tdTN-cMPK1 using the *Agrobacterium*-based method. T1 transformants selected by hygromycin only or by hygromycin and chlorsulfuron together, respectively, were genotyped by PCR with the primer set MpMPK1_-549F and MpMPK1_g540R, and the amplified fragments were directly sequenced. G1 gemmae of the pMpGWB337tdTN-cMPK1-harboring genome-edited lines were further confirmed to have the same mutations as those in the respective parental T1 lines. G2 gemmae of line #4 were planted on a 9-cm plastic plate containing half-strength B5 medium with 1 µM dexamethasone (DEX) and further treated with a drop of 1 µM DEX. The plate was incubated in a 3722°C air incubator for 80 min and then moved to a 22°C growth room. In 24 h, the same procedures were repeated, and the plate was incubated in the 22°C growth room for 13 days. Observation of tdTomato fluorescence was performed using a stereoscope (M205C, Leica) with the Ds-Red2 filter set (Leica). Induction of cDNA deletion was examined by genomic PCR using the primer sets 1 (MpEF-P_seqL1 and MpMPK1_c1131R_STP), 2 (MpEF-P_seqL1 and tdTomato_753R), and 3 (MpMPK1_-549F and MpMPK1_g1010R). The primers used are listed in Supplementary Table S1.

## Results

### Improved genome editing efficiency in *M. polymorpha* by using Arabidopsis codon optimized Cas9

In the previous report, we co-introduced a construct for gRNA expression with a 2 kbp Mp*U6-1* promoter and another for the expression of human-codon-optimized Cas9-NLS (*hCas9-NLS*) to reconstruct a CRISPR/Cas9 system in *M. polymorpha*, and succeeded in obtaining plants whose target locus was edited [25]. However, the genome editing efficiency was very low, under 0.5%; that is, only a few plants with genome editing events were obtained for each co-transformation of sporelings derived from two sporangia. To improve the genome editing efficiency to a practical level, we examined the effects of codon optimization by replacing *hCas9-NLS* with *Atco-Cas9-NLS* (although the *35S* terminator was also replaced by the *Pisum sativum rbcS3A* terminator [12], this difference seems negligible (unpublished data)). In this new Cas9 expression plasmid, designated pMpGE006, *Atco-Cas9* was driven by the constitutive Mp*EF*_*pro*_ promoter, which is preferentially expressed in meristematic tissues in *M. polymorpha* (Fig. 1A) [32]. For the expression of a gRNA, the same gRNA expression vector used in the previous study was again utilized [25].

To evaluate genome-editing efficiencies, we firstly chose the Mp*ARF1* (*AUXIN RESPONSE FACTOR1*) transcription factor as a target, which had been used in the previous report [25, 33]. Since Mp*arf1* mutants are known to show an NAA resistant phenotype [34, 35], auxin-resistant T1 transformants with mutations in the Mp*ARF1* locus can be positively selected. The gRNA expression plasmid for Mp*ARF1*, pMpGWB301_ARF1_1, which had previously been proved to be effective [25], was also used in this experiment (Figs. 1A, S2). A single co-transformation of sporelings with pMpGE006 and pMpGWB301_ARF1_1 yielded hundreds of NAA-resistant plants (Figs. 1B, S3), whereas the same experiment with the *hCas9* plasmid,instead of pMpGE006, yielded few NAA-resistant lines (Fig. S3). We analyzed the target sequence of Mp*ARF1* in 16 NAA-resistant T1 co-transformants of pMpGE006 and pMpGWB301_ARF1_1 by direct sequencing, all of which harbored indels and/or base substitutions in the target site (Fig. 1C). Therefore, we conclude that the Atco-Cas9 product has much higher efficiency than hCas9.

Next, we examined whether mutants could be isolated without the NAA-based phenotypic selection. T1 co-transformants were selected only by using hygromycin and chlorsulfuron, which are selection markers for pMpGE006 and pMpGWB301_ARF1_1, respectively. Direct sequencing analyses showed that 75% of the randomly selected T1 plants (12 of 16) had some mutations in the target sequence of Mp*ARF1* (Fig. 1D, E). Although one plant showed a mosaic sequence pattern (Fig. 1D, E), the majority was non-mosaic, suggesting that the genome editing events had occurred in an early phase of transformation. From these data, we concluded that the efficiency of genome editing using the Atco-Cas9 expression cassette is high enough to isolate mutants by direct sequencing analysis of target genes without any target-gene-dependent phenotypic selections.

### Construction of genome editing vector series in *M. polymorpha*

In addition to pMpGE006, we constructed a series of genome editing vectors for addressing the necessity of various applications of genome editing. We constructed a single binary vector system with both gRNA and Cas9 expression cassettes to make more easy-to-handle vectors (Fig. 2). pMpGE010 and pMpGE011 are based on the Gateway cloning system (Fig. 2A). The entry clone vector harbors gRNA expression cassettes in pENTR/D-TOPO, designated as pMpGE_En01 to pMpGE_En03 (Fig. 2A). pMpGE_En01 contains a 2 kbp Mp*U6-1_pro_* for gRNA expression and the cloning site for double stranded oligos with a guide sequence using the In-Fusion reaction (Fig. S1A). pMpGE_En02 contains a 500 bp Mp*U6-1_pro_* sequence for gRNA expression and is designed for restriction enzyme-based cloning (Fig. S1B). This vector does not contain a purine nucleotide at the 5’ end of the gRNA cloning site, which is used to start transcription by RNA polymerase III [36]. pMpGE_En03 is identical to pMpGE_En02 except for a built-in “extra initial G” (Figs. 2A, S1B). The “extra initial G” also exists in pMpGE_En01 (Fig. S1A). For binary vectors, pMpGE010 was constructed using pMpGWB103 [18] as a backbone vector, which harbors a hygromycin resistance cassette. Likewise, pMpGE011 was constructed using pMpGWB303, which harbors a chlorsulfuron resistance cassette.

**Fig 2.**
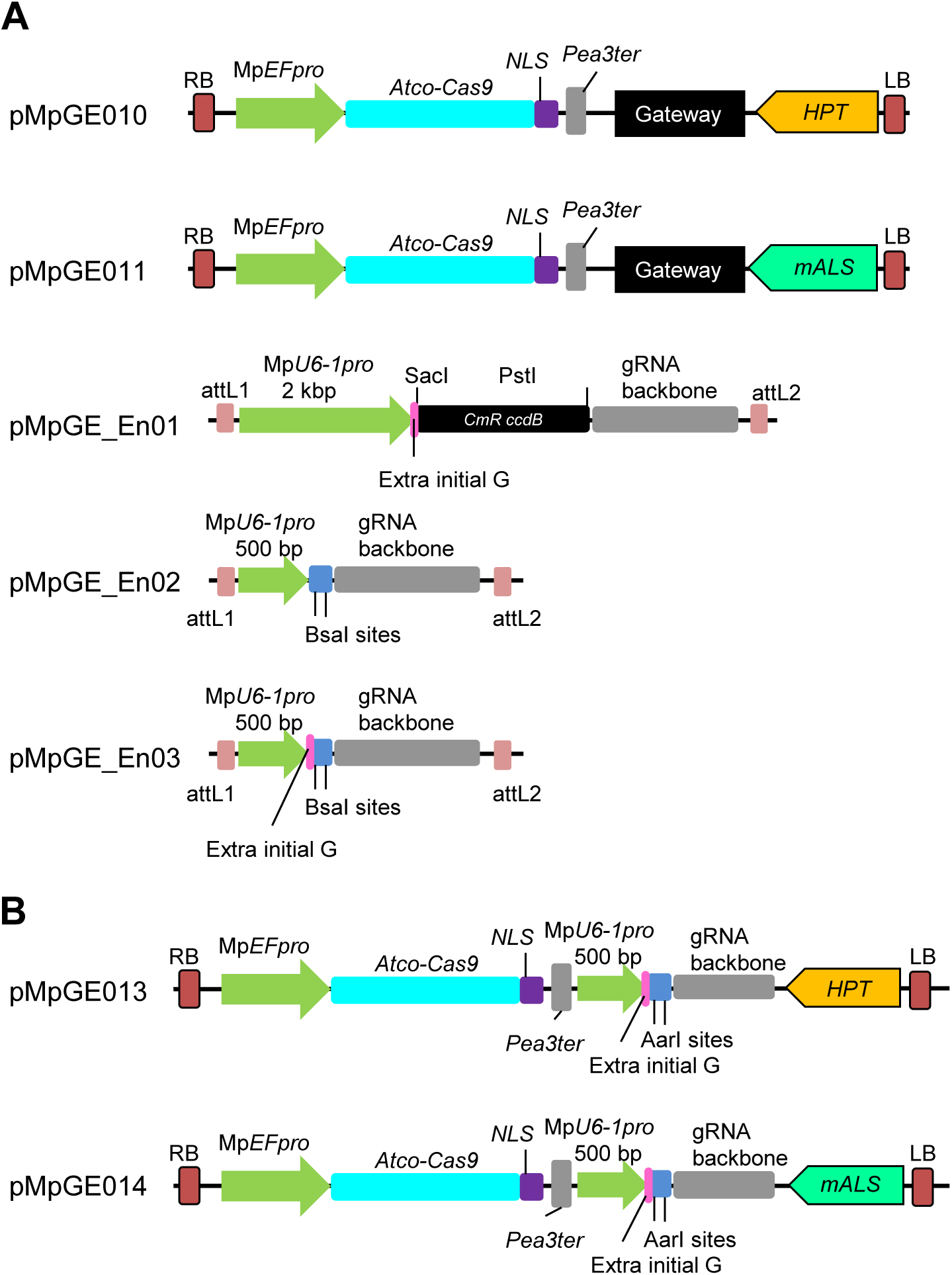
All-in-one vector systems for genome editing in *M. polymorpha*. (A) Designs of Gateway-based all-in-one binary vectors and entry plasmids for gRNA cloning. pMpGE010 and pMpGE011 contain a cassette for the expression of Atco-Cas9 fused with an NLS under the control of Mp*EF*_*pro*_, a Gateway cassette, and a cassette for the expression of hygromycin phosphotransferase (*HPT*) in pMpGE010 and mutated acetolactate synthase (*mALS*) in pMpGE011. pMpGE_En01 contains recognition sites for two restriction enzymes, SacI and PstI, upstream of a gRNA backbone for the insertion of a guide sequence by In-Fusion/Gibson cloning, which automatically places a G nucleotide for transcription initiation by RNA polymerase III (extra initial G). Expression of single guide RNAs is controlled by a 2 kbp fragment of Mp*U6-1_pro_*. pMpGE_En02 and pMpGE_En03 contain two BsaI recognition sites upstream of the gRNA backbone for the insertion of a guide sequence by ligation without or with an “extra initial G,” whose expression is under the control of a 500 bp Mp*U6-1_pro_* fragment. For all the entry vectors, the gRNA cassette is flanked by the attL1 and attL2 sequences and is thus transferrable to the Gateway cassette in pMpGE010 or pMpGWB011 by the LR reaction. (B) Designs of all-in-one binary vectors for direct gRNA cloning. pMpGE013 (*HPT* marker) and pMpGE014 (*mALS* marker) contain the Atco-Cas9-NLS expression cassette, a unique AarI site in the upstream of the gRNA backbone for insertion of a guide sequence by ligation with an “extra initial G,” whose expression is under the control of a 500 bp Mp*U6-1_pro_* fragment.

For easier construction, binary vectors based on restriction-enzyme-based cloning, pMpGE013 and pMpGE014, were also constructed using pMpGE010 and pMpGE011 as backbones, respectively. Double stranded oligos harboring the target sequence of gRNAs can be directly cloned into pMpGE013 and pMpGE014, which contain the “extra initial G,” at the AarI restriction enzyme sites (Fig. 2B). These vectors provide a low-cost alternative for gRNA cloning.

### Evaluation of the efficiency of genome editing vectors

To evaluate genome-editing efficiencies in the new system, we exploited targeted mutagenesis at two loci (Fig. 3). We cloned into pMpGE_En01 a gRNA harboring a guide sequence identical to that of pMpGWB301_ARF1_1 and transferred it to pMpGE010 (Fig. S2). The constructed vector was used for transformation of sporelings and the transformants were selected with hygromycin only on auxin-free media. Direct sequencing of the ARF1_1 gRNA target site revealed that 75% of T1 plants (24 of 32) had mutations at the target site (Fig. 3A, B). A similar result was obtained with pMpGE011 (Fig. S4A). Thus, the efficiency of targeted mutagenesis with the single vector system was comparable to that with the double vector system.

**Fig 3.**
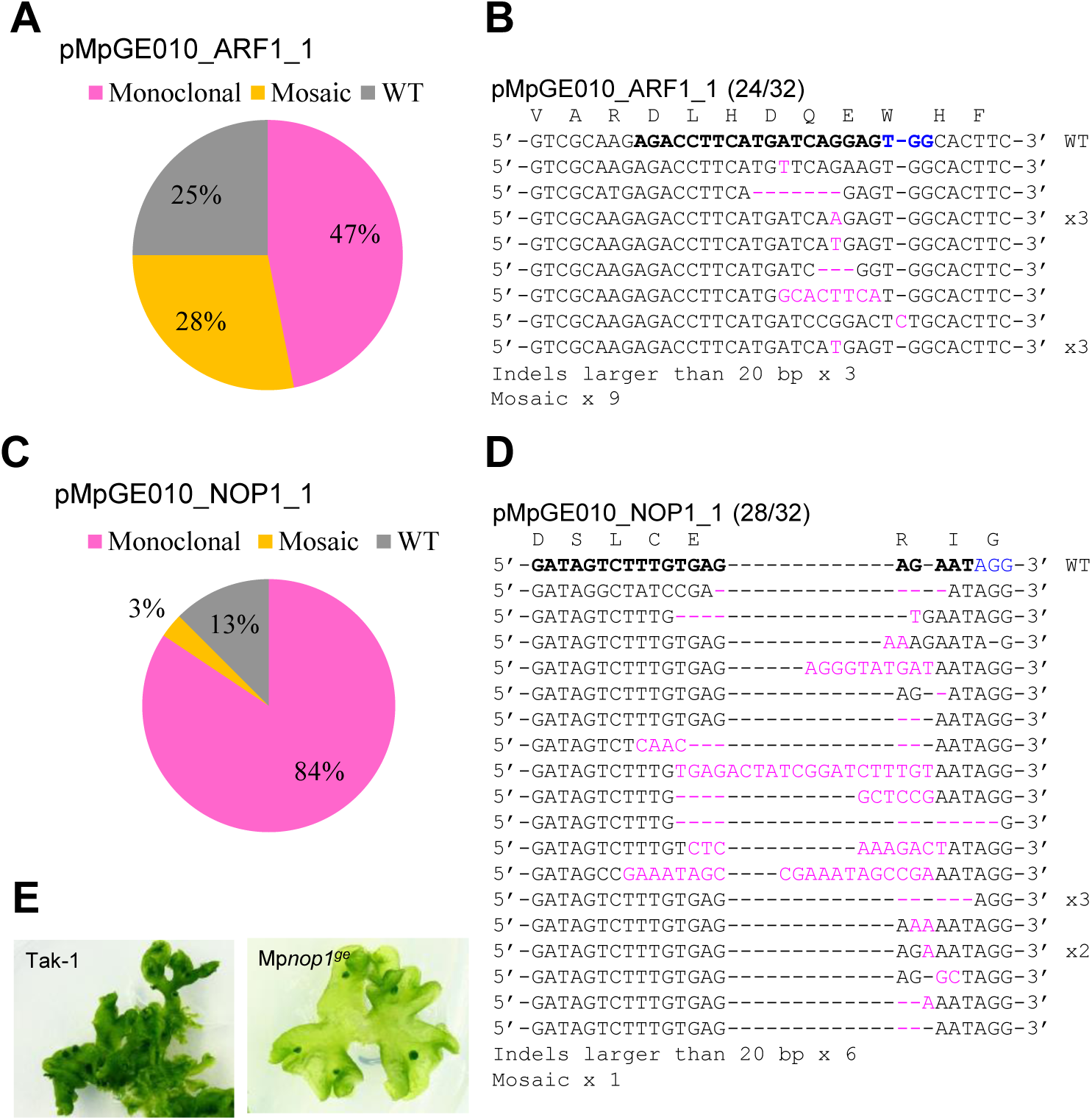
High-efficiency genome editing by the gateway-based system in *M. polymorpha*. Sporelings were transformed with pMpGE010 harboring ARF1_1 gRNA (A and B) or NOP1_1 gRNA (C-E). Randomly selected transformants were genotyped. Proportions of genome editing patterns (A and C) and sequences of the target sites (B and D) in 32 T1 transformants are shown as in Fig. 1. Panel E shows photographs of a wild-type plant (Tak-1) and a T1 transformant that exhibited the typical *NOP1*-defective “transparent” phenotype (Mp*nop1^ge^*).

To examine whether genome editing could be applied at other loci, we chose a gene, *NOPPERABO1* (Mp*NOP1*), which encodes a plant U-box E3 ubiquitin ligase that is responsible for air-chamber formation in *M. polymorpha* [37]. Since Mp*nop1* mutants form transparent thalli due to the lack of air chambers, they can be easily distinguished from wild-type plants with the naked eye. An Mp*NOP1*-targeting gRNA, NOP1_1 (Fig. S2), was introduced into sporelings with pMpGE010. Direct sequencing analysis revealed that 87.5% of T1 plants (28 of 32) had some mutation in the target sequence (Fig. 3C, D). Many of the mutant lines exhibited the transparent phenotype in the entire body, reflecting the frequency of non-mosaic sequence reads (Fig. 3E).

We also assessed a shorter (0.5 kbp) Mp*U6-1* promoter by comparing pMpGE_En01 and pMpGE_En02. Genome-editing efficiency with pMpGE_En02 was comparable to that with pMpGE_En01, when examined with the gRNA NOP1_1 (Fig. S4B). This result suggests that the 0.5 kbp Mp*U6-1* promoter is sufficient to drive gRNA expression for efficient genome editing.

### Influence of gRNA lengths to genome editing efficiency

Next, we assessed the length of the gRNA guide sequence. It has previously been reported that truncated gRNA guide sequences (17 nt and 18 nt) show low off-target activities in mammalian cells [38]. We chose a guide sequence (5’-CAAACCGGAATGAGTCAGCT-3’), which targets the exon of the Mp*NOP1* gene (NOP1_2; Fig. S2) and cloned various lengths (20 nt, 18 nt, 17 nt, and 16 nt) of the sequence into pMpGE_En02, which does not have the “extra initial G.” Genome-editing efficiencies were scored by classifying the penetrance of the transparent phenotype due to mutations in the Mp*NOP1* gene: class I, plants with the transparent phenotype observed throughout; class II, plants with the transparent phenotype observed in a mosaic fashion; and class III, plants with no obvious phenotype (Fig. 4A). The results clearly showed that guide sequences of 17 nt or fewer had much lower genome-editing efficiencies than those of 18 nt or more (Fig. 4B).

**Fig 4.**
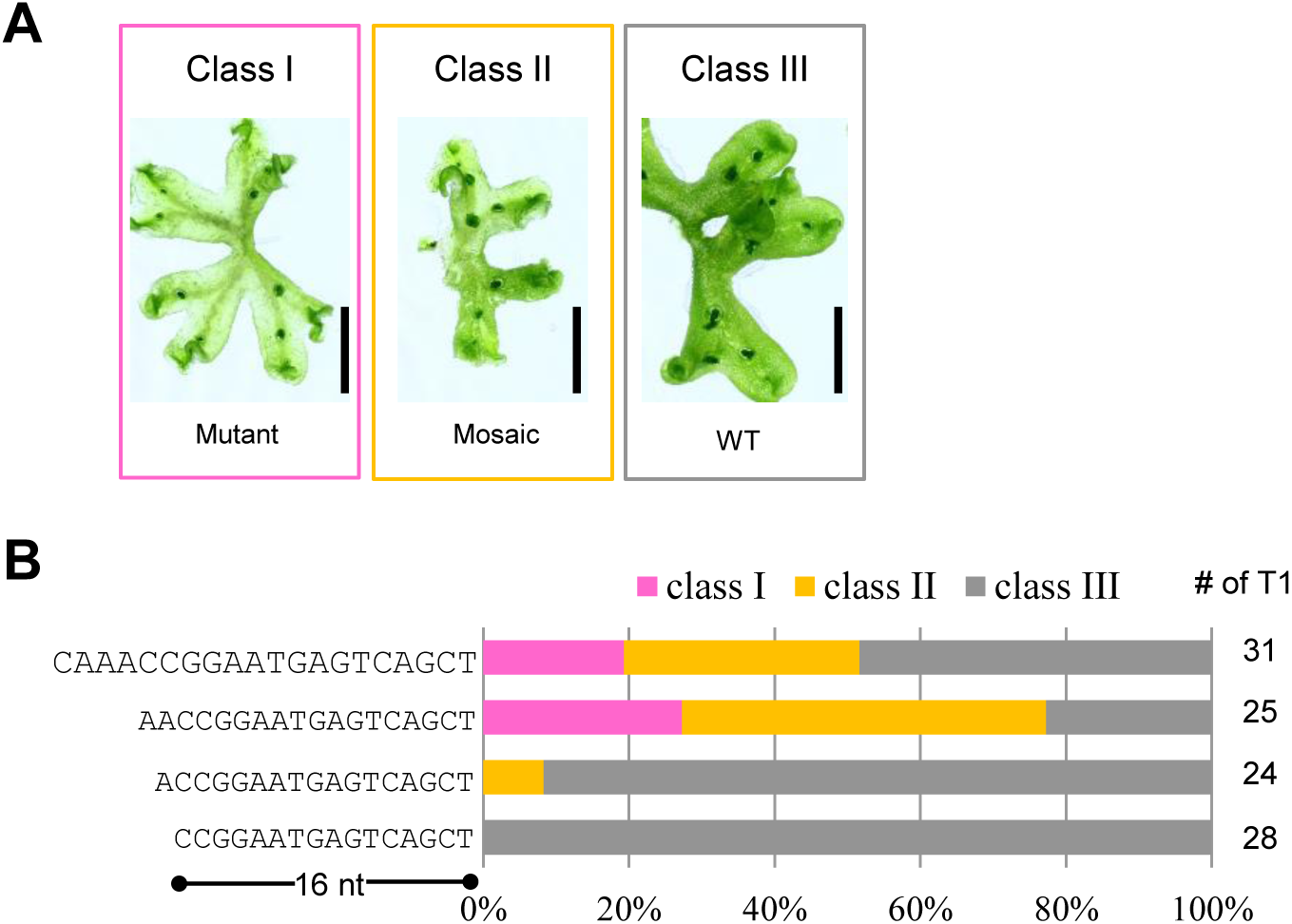
Effects of guide sequence lengths (A) Classification of phenotypes in Mp*NOP1* genome editing lines by the appearance patterns of transparent portions. Class I, entirely transparent (Mutant); class II, mosaic of transparent and non-transparent sectors (Mosaic); class III, entirely non-transparent (wild-type, WT). Scale bars = 1 cm. (B) Proportions of mutant phenotype classes in T1 plants transformed with Mp*NOP1*-targeting gRNAs (NOP1_2) of different lengths. The shortest gRNA guide sequence tested was 16 nt. The numbers of T1 plants inspected are shown on the right hand side.

The low genome editing efficiency when using the gRNAs with 17-nt and 16-nt guide lengths could have been caused by their reduced expression levels due to the lack of an initial guanine. Thus, we investigated the effects of addition of an guanine to the 5’ end of gRNAs (using pMpGE_En03), which should facilitate transcription by pol III [39]. However, no clear improvement in genome editing efficiency was observed (Fig. S5). These results suggest that the occurrence of lower genome editing events in *M. polymorpha* may strongly depend on the length of a guide sequence that perfectly matches the target genome, which should be 18 nt or longer.

### Assessments of off target effects

As plants obtained by transformation with pMpGE010/011 stably express Cas9 and a gRNA, genome sites with sequences similar to that of a gRNA are always at risk of genome editing. Sequencing analysis of the genome of more than 30 T1 plants that harbored on-target mutations in the ARF1_1 or NOP1_1 target locus revealed no mutation at any of the three most potential off-target sites (Table 1). Collectively, it is suggested that genome editing efficiencies in off-target sites are much lower than those in on-target sites in *M. polymorpha*.

**Table 1:**
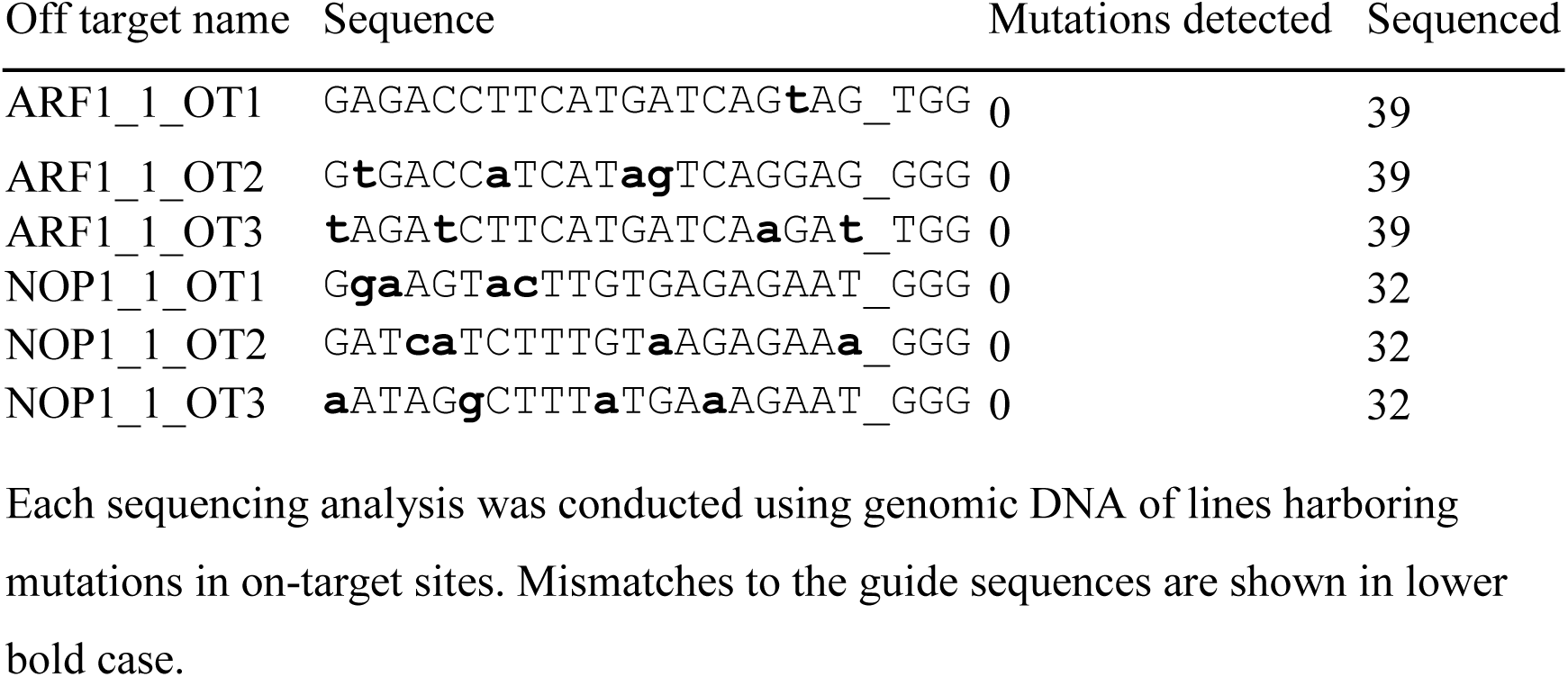
Off target analysis of T1 transformants. Each sequencing analysis was conducted using genomic DNA of lines harboring mutations in on-target sites. Mismatches to the guide sequences are shown in lower bold case.

### *De novo* mutations after prolonged culture

The stable expression of the CRISPR/Cas9 system should provide continuous opportunities for targeted mutagenesis in transformants until a mutation has been introduced. We analyzed gemmae (G1 generation [18]) derived from pMpGE010_NOP1_1 T1 transformants that had shown sectors of transparent (mutant) and non-transparent (wild-type) thallus regions in one individual (Fig. 5A). As expected, G1 gammalings obtained from transparent sectors basically inherited the same mutations as those found in their corresponding T1 sectors (Fig. 5B).Concurrently, some gammalings from non-transparent sectors were found to contain *de novo* mutations, which were different from those in the transparent sectors of the same parent individuals (Fig. 5B, C). These results indicate that various allelic mutants can be isolated from single non-mutated transformants.

**Fig 5.**
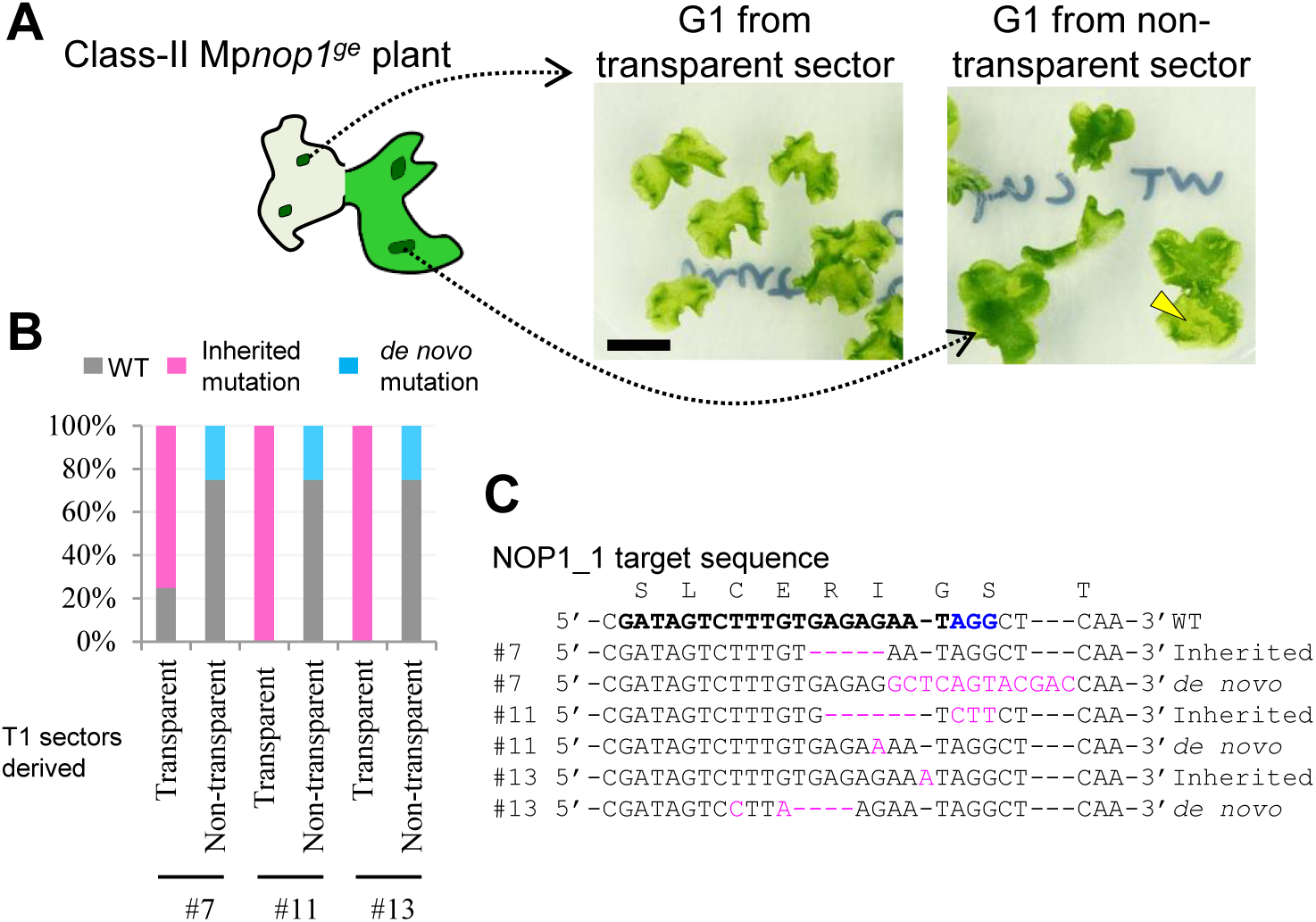
*De novo* mutations found in the G1 generation (A) Delayed generation of new mutants from non-mutated sectors. G1 gemmae formed on a transparent or non-transparent sector of a class-II Mp*nop1^ge^* plant, described in Fig. 4, were grown. Arrowhead shows a transparent sector due to a *de novo* mutation. Scale bar = 1 cm. (B) Proportions of genotypes in G1 populations derived from three independent class-II Mp*nop1^ge^* T1 lines (#7, #11, and #13). “Inherited mutation” and “*de novo* mutation” indicate mutations identical to or different from those identified in individual T1 lines, respectively. (C) Target-site sequences in lines with “Inherited mutation” and “*de novo* mutation” by direct sequencing analysis. Inserted or substituted bases are colored in magenta. The target guide sequence of NOP1_1 is shown in bold face with the PAM sequence in blue.

### Induction of large deletion using two gRNAs

Previous studies have reported that CRISPR/Cas9 system could induce deletions between two gRNA target sites in mosses [8]. Accordingly, induction of large deletions using two gRNAs was tested in *M. polymorpha*. We designed four gRNAs to the Mp*NOP1* gene, NOP1_3, NOP1_4, NOP1_5, and NOP1_6 (Fig. 6A). Simultaneous introduction of pMpGE013 and pMpGE014 harboring different combinations of gRNAs was conducted, and transformants were selected with both hygromycin and chlorsulfuron. Deletions of expected sizes from the gRNA combinations, that is, a 0.5 kbp deletion with NOP1_3 and NOP1_4, a 1.5 kbp deletion with NOP1_3 and NOP1_5, and a 4.5 kbp deletion with NOP1_3 and NOP1_6, were detected, respectively (Fig. 6B). The efficiencies of induction of large deletions were almost comparable regardless of the deletion size: 9/26 for 0.5 kbp,4/20 for 1.5 kbp, and 6/23 for 4.5 kbp (Fig. 6C). These large-deletion lines displayed the Mp*nop1* phenotype. (Fig. S6). Collectively, the induction of large deletions with this system would also be applicable for functional analysis of genes in general in *M. polymorpha*.

**Fig 6.**
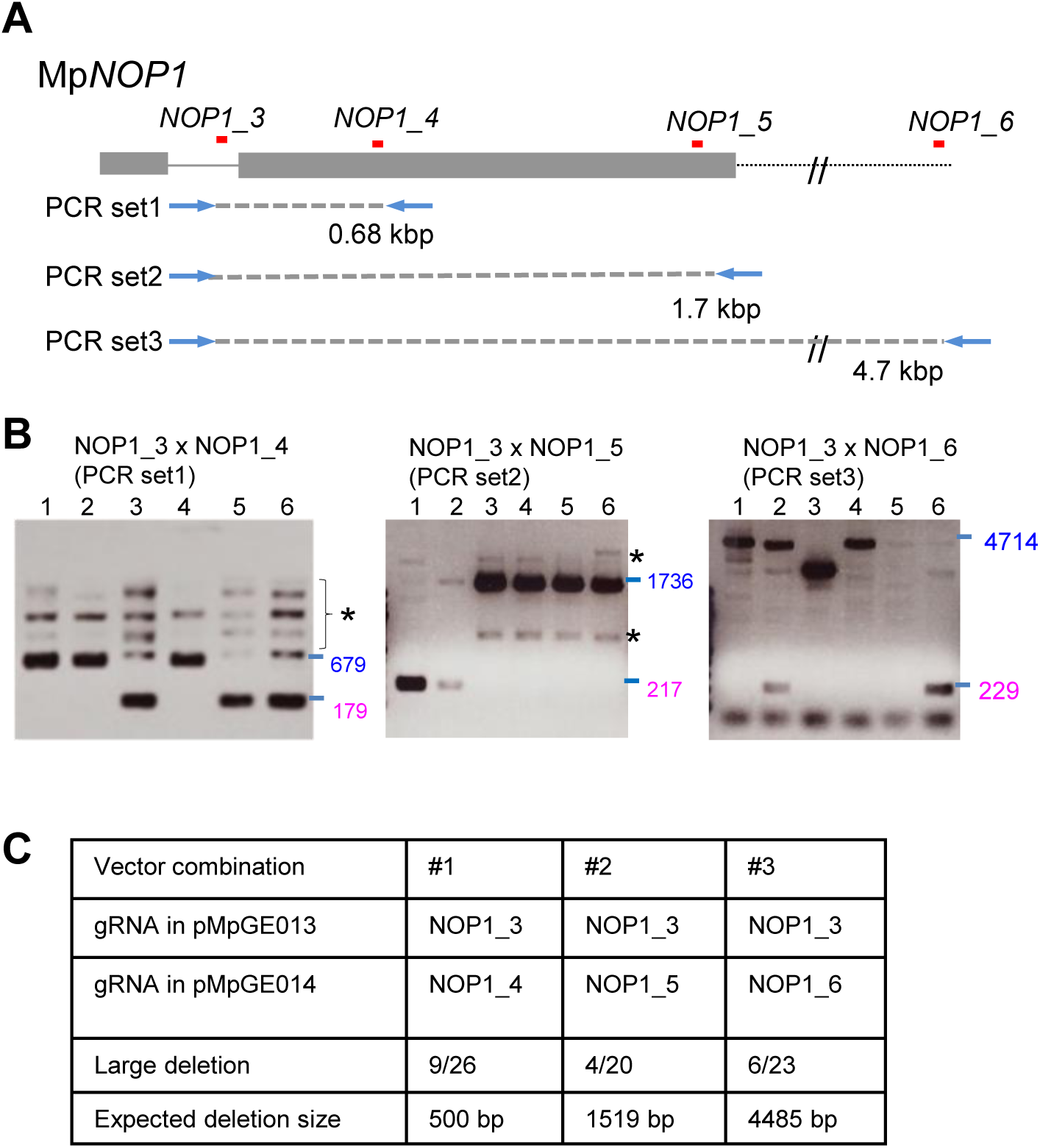
Induction of large deletions using co-transformation of two genome editing vectors (A) Design of gRNAs to the Mp*NOP1* locus to dissect efficiencies of induction of large deletions. Grey boxes and lines indicate exons and introns. The dotted line indicates the downstream region of Mp*NOP1*. gRNA positions are shown in red lines. Primer sets for PCR-based genotyping are also shown. (B) Representative images of electrophoresis of PCR-based genotyping. The gRNAs and primer sets used in the genotyping are shown. Expected sizes of the PCR products from the wild-type genome are colored in blue. Expected sizes of the PCR products from the genome in which inductions of large deletion occurred are colored in magenta. PCR products from non-specific amplifications are indicated by asterisks. (C) Summary table of the co-transformation experiment using pMpGE013 and pMpGE014. The ratio in the “Large deletion” row shows the number of T1 plants harboring the expected large deletion out of all the T1 plants inspected.

### One-step generation of conditional knockout mutants

Mutants that exhibit phenotypes under certain conditions are useful when a gene of interest has multiple roles in the life cycle and/or, in particular, essential functions. In this study, we provide methodology for obtaining conditional knockout mutants in a single step using the CRISPR/Cas9 system for mutagenesis and an inducible Cre-*lox*P site-specific recombination for complementation. As our model, we chose one of the three mitogen-activated protein kinase (MAPK) genes in *M. polymorpha*, Mp*MPK1* [16], which was predicted to be an essential gene because attempts to obtain knockout mutants by the homologous-recombination-based gene targeting method [22] had failed.

We constructed pMpGE010 harboring a gRNA that was designed to target the first exon-intron junction in Mp*MPK1* (Fig. 7A). Because the conjunction between exons 1 and 2 does not reconstitute the PAM sequence for this gRNA (Fig. 7A), it is not supposed to target the Mp*MPK1* cDNA. Thus, for complementation, we cloned an Mp*MPK1* cDNA into the binary vector pMpGWB337 [23] or its derivative pMpGWB337tdTN (Fig. S7), both of which normally drive expression of the cDNA but inducibly allow deletion of the cDNA and expression of a fluorescent protein after application of heat shock and DEX. Transformation of sporelings with only the Mp*MPK1* CRISPR vector yielded a small number of mosaic mutants but no monoclonal frameshift mutants (Fig. 7B), which is indicative of possible lethality for genome editing at the target site. However, simultaneous transformation with the same CRISPR vector and either of the complementation vectors described above gave rise to monoclonal frameshift mutants at high frequencies (Fig. 7B, C). These data suggest that the cDNA resistant to the gRNA allowed complementation of deleterious mutations in the endogenous Mp*MPK1* locus.

**Fig 7.**
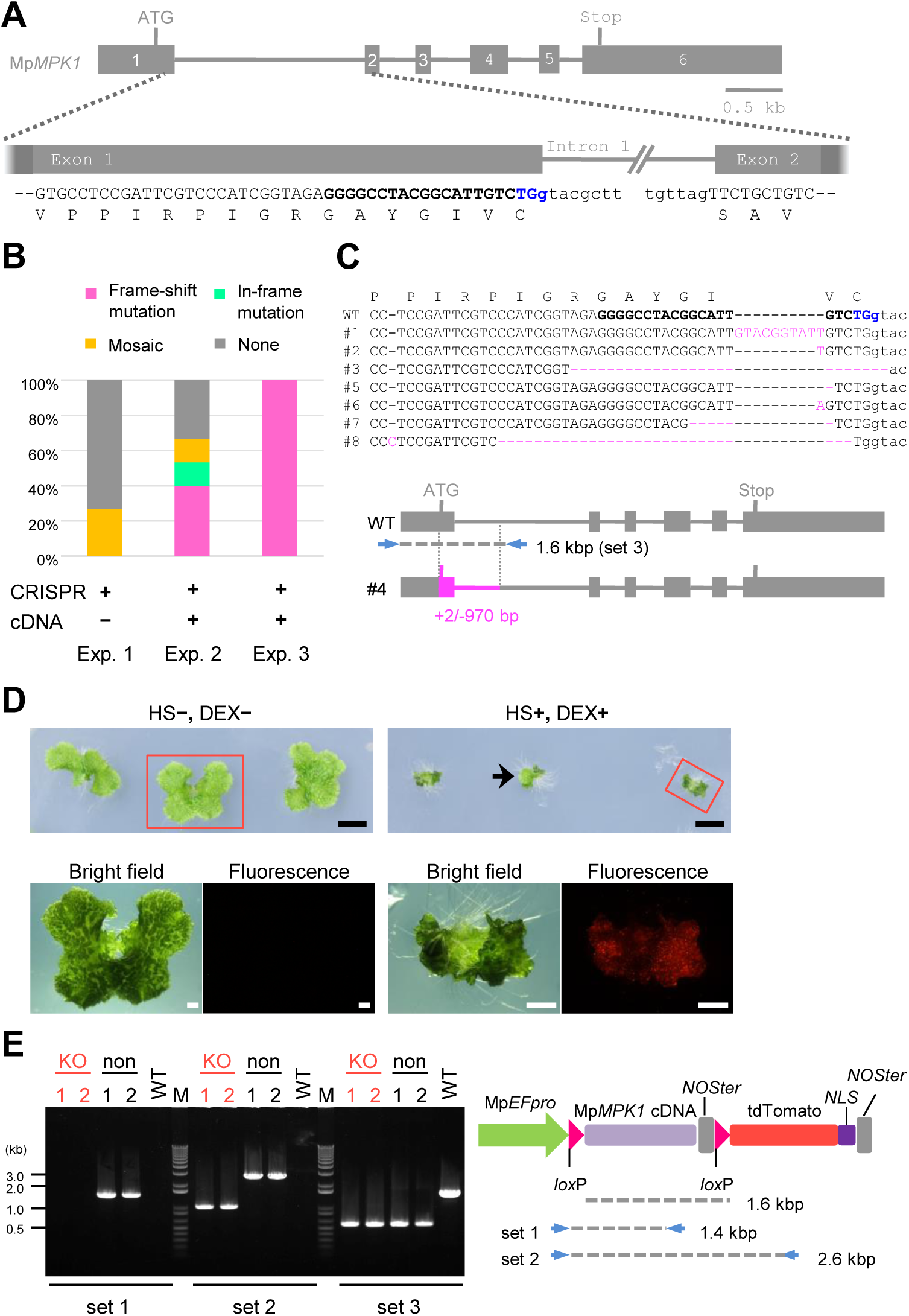
Generation of conditional knockout mutants of an essential gene (A) Gene model of the Mp*MPK1* gene and the gRNA target site. Grey boxes and lines indicate exons and introns. “ATG” and “Stop” denote the predicated initiation and termination codons. PAM, target, and intron sequences are shown in blue, in bold, and by small letters, respectively. Note that the third base (g) of the PAM sequence is in intron 1 and that the first base of exon 2 is not G. (B) Targeted mutagenesis rate of the endogenous Mp*MPK1* locus by CRISPR/Cas9. Bar graphs show proportions of clonal frameshift (magenta), clonal in-frame (green), mosaic (yellow), and no mutations (grey) that were identified in plants transformed with pMpGE010 containing the gRNA shown in (A) only (Exp. 1) or together with Mp*MPK1* cDNA-containing pMpGWB337 (Exp. 2) or pMpGWB337tdTN (Exp. 3). (C) Target-site sequences of plants obtained in Exp. 3. Sequences of wild type (WT) and transgenic lines (#1 to #8 except #4) are shown as in (A). Inserted or deleted bases are colored in magenta with their numbers in parentheses. In line #4, a 2 bp insertion and a large 970 bp deletion that covers the predicted initiation codon were identified (magenta). (D) Growth defect manifested by conditional deletion of the Mp*MPK1* cDNA. Gemmae of transgenic line #4 were (+) or were not (–) subjected to heat shock (HS) and dexamethasone (DEX) treatment on day 0 and day 1 and grown at 22°C for 2 weeks (top) before observation by fluorescent microscopy for tdTomato-NLS (bottom).Scale bars = 5 mm (top); 1 mm (bottom). (E) Confirmation of deletion events by genomic PCR. From HS/DEX-treated plants (line #4), tdTomato-positive, disorganized tissues (“KO”) and tdTomato-negative, thallus-looking sectors [such as the arrow in (D); “non"] were collected from two different individuals and analyzed by genomic PCR using primer sets 1 and 2 shown in the schematic illustration of the construct used (predicted product sizes are indicated) and primer set 3 shown in panel (C).

These complemented mutants grew normally and were fluorescent negative under mock conditions, whereas upon heat shock and DEX treatment, they grew extremely slowly and became fluorescent positive (Fig. 7D). Induction of cDNA deletion was confirmed by genomic PCR analysis (Fig. 7E). These results suggest that Mp*MPK1* is indeed a nearly essential gene and, more importantly, demonstrated that conditional knockout mutants of an essential gene can be generated by a simple procedure using a CRISPR/Cas9 vector and a pMpGWB337 derivative in *M. polymorpha*.

## Discussion

Using the *Atco-Cas9* expression cassette, we successfully optimized the CRISPR/Cas9-based genome editing system for *M. polymorpha* and improved the efficiency to a degree that does not require target gene-based phenotypic selection. This is consistent with a previous report that codon optimization of Cas9 lead to significant improvement in efficiency [40]. In the pMpGE010/011 system, over 70% of transformants underwent targeted mutagenesis, shown by the two gRNAs ARF1_1 and NOP1_1 (Figs. 1, 2). These results indicate that the pMpGE010/011 vectors are highly reliable for obtaining genome-edited lines in *M. polymorpha*. Since the GC content of the *M. polymorpha* genome is 49.8% [16], Cas9 target sites with NGG PAM sequences can be found at high probability, and the selection of gRNA target sites with small numbers of off-target sites is possible. Taken together, we conclude that the pMpGE010/pMpGE011 system is feasible to use for functional genetics in *M. polymorpha* with likely avoidance of off-target effects.

As the system presented here allows constant expression of both Cas9 and a gRNA, two risks are conceivable: (i) a side effect of the overexpression of Cas9 protein on plant development and (ii) genome editing events at off-target sites.However, both risks appear to be negligible. Firstly, Mp*nop1* mutants obtained with our CRISPR system were indistinguishable in terms of growth and morphology from those obtained by T-DNA tagging [22] or gene targeting [37] (Fig. 3A). In addition, the conditional Mp*mpk1* mutants generated in this study did not show any noticeable developmental defects (Fig. 7D). These observations suggest that there is no detrimental effect of Cas9 overexpression on plant development.

Sequencing of several potential off-target sites of 20-nt guide sequences revealed remarkably low off-target effects: no mutation was observed at any of the sites in over 30 lines that had on-target mutations. In plants, a similar low-level off-target effect was reported for Arabidopsis [14]. It was reported that truncated gRNA guide sequences (17 nt and 18 nt) show low off-target activities in mammalian cells [38]. In *M. polymorpha*, 17-nt and shorter gRNA guide sequences were not effective (Fig. 3). Taken together with the observed low off-target effects, to avoid unstable outcomes in isolating genome-edited lines, it is recommended to use gRNAs with 18-nt or longer guide sequences in *M. polymorpha*.

Constant expression of the CRISPR system allows transformants that have not had any alteration to potentially acquire mutations later at the target site in random cells during growth. Indeed, we observed that pMpGE010-transformed thalli with no target mutation in the T1 generation produced gammae with *de novo* mutations (Fig. 5). This feature would be convenient for isolating mutant alleles without further transformation and could also be used for mosaic analyses. Conversely, even if a monoclonal mutant pattern is detected in genotyping using a portion of T1 plants, it does not guarantee that the whole plant has the same genotype. For genotyping, it is recommended to use a piece of T1 tissue from the basal side of the apical notch, as G1 gemmae derived from its apical-side tissues are most likely to be clones, with an identical genotype to that found in T1 [17].

Diploid or polyploid plants can bear lethal mutations heterozygously if recessive, whereas *M. polymorpha*, a haploid-dominant plant, cannot. Therefore, functional analyses of essential genes require alternate strategies. Flores-Sandoval (2016) reported an inducible system for artificial microRNA expression in *M. polymorpha*. Another strategy would be to create conditional knockout mutants. In mice, this is usually achieved by inserting *lox*P sites into two different introns in the same direction with homologous-recombination-mediated gene targeting, and then expressing Cre recombinase at a specific time and/or location to remove an essential exon [41]. For *M. polymorpha*, we avoided using a laborious gene-targeting-based strategy. Instead, we established a method to generate a conditional knockout mutant by simultaneous introduction of a mutation in the endogenous target gene by the CRISPR/Cas9 system and of a conditionally removable complementation gene as a transgene.

In this strategy, the complementation gene cassette must have a structure that cannot be targeted by the gRNA used for knocking out the target gene. Although this “gRNA-resistant” complementation cassette can be prepared by introducing synonymous substitutions in the matching sequence, extra time is required. Thus, if possible, we recommend complementation, by expressing a non-modified cDNA, of mutations caused by a non-cDNA-targeting gRNA, which can be designed at exon-intron junctions. A DNA fragment for complementation can be inserted between two *lox*P sites in pMpGWB337 [23] or its derivatives with different fluorescent protein markers (Fig. S7), all-in-one vectors equipped with a floxed Gateway cassette for introducing a complementing gene and with a heat-shock- and DEX-inducible Cre recombinase expression cassette.

Using this method, we successfully isolated transformants with a mutation in the Mp*MPK1* locus, which was expected to be essential. Mp*MPK1* is one of the three genes encoding canonical MAPKs and is most closely related to those categorized as groups A and B for plant MAPKs [16, 42]. Major Arabidopsis MAPKs in these groups (MPK3/MPK6 and MPK4, respectively) have been shown to regulate various aspects of growth and development [43]: *mpk3 mpk6* double mutants are embryonic lethal [44], and *mpk4* mutants show growth retardation with a cytokinesis defect [45]. The co-transformation-generated transgenic plants that had a mutation in the endogenous Mp*MPK1* locus exhibited no growth defect due to complementation by the expression of an Mp*MPK1* cDNA (Fig. 7). Induction of Cre-*lox*P recombination by heat shock and DEX treatment resulted in severe growth defects, indicating that MpMPK1 plays a critical role in the regulation of growth and development in *M. polymorpha* and that the conditional induction of the mutant phenotypes successfully occurred. Sectors of non-abnormal tissues sometimes arose, probably due to occasional failure of the Cre-*lox*P recombination. Thus, it is important to select conditional lines with highly efficient recombination induction from independently isolated candidates. The conditional knockout system developed in this study should not only facilitate functional analyses of essential genes, but also be useful for uncovering functions of non-essential genes in specific locations or timings.

DSBs created by CRISPR/Cas9-based genome editing are usually repaired by the NHEJ pathway. Although the error-prone NHEJ repair pathway randomly inserts or deletes bases at the DSB sites, some tendencies were observed in *M. polymorpha*. For example, mutants with a >20 bp deletion were frequently obtained (Figs. 1, 2), while such mutations are rare in genome editing in Arabidopsis [12, 46]. This suggests that the NHEJ-based repair activity in *M. polymorpha* is relatively weaker than that in Arabidopsis. Consistent with this idea, homologous recombination-based gene targeting is possible in *M. polymorpha* with the average rate of 2-3%, which is much higher than in Arabidopsis [22]. Since DSBs induced by the CRISPR/Cas9 system were reported to increase the efficiency of homologous recombination [47], it would be possible to improve the gene targeting efficiency in *M. polymorpha* by combination with the vectors presented in this study. Another frequently observed feature was the repair of DSBs by the microhomology-mediated end joining (MMEJ) pathway (Fig. S8). Thus, precise integration of a DNA construct into target loci assisted by MMEJ, which is available for animals [48, 49], would also be possible in *M. polymorpha*.

Construction of the genome editing vectors in the present study was very simple and inexpensive. Together with the fact that *M. polymorpha* has low redundancy and small numbers of regulatory genes, our highly efficient CRISPR/Cas9 system would allow genome-editing-based genetic screens targeting all genes in a large subset of the genome, e.g., protein kinases, transcription factors, and miRNAs [16, 50, 51]. Since the “pMpGE” CRISPR/Cas9-based genome editing vectors reported in this study possess 35S promoter-based marker cassettes, the vectors can be utilized for genome editing in other plants in which Mp*EF*_*pro*_ and Mp*U6*_*pro*_ are operable. Highly efficient genome editing should facilitate uncovering gene regulatory networks that evolved for the land adaptation of plants and that underlie subsequent successful expansion of land plants.

## Supporting information

S1 Supplementary Figures S1-S8 (PDF)

S2 Supplementary Table S1(.xlsx file)

## Data availability

The complete nucleotide sequences of the pMpGEs have been deposited in the GenBank/EMBL/DDBJ databases under the accessions nos. LC090754 to LC090757. pMpGE plasmids can be obtained from Addgene (www.addgene.org; plasmid numbers 71534–71537).

## Acknowledgements

We thank H. Puchta for the gifts of the pDe-CAS9 material, including Atco-Cas9. We thank S. Yamaoka, K.T. Yamato, S. Zachgo, Y. Osakabe, M. Endo, and S. Toki for helpful discussion. We also thank J. Haseloff and B. Pollak for the gRNA sequence of the Mp*NOP1* gene. We thank M. Fukuhara, Y. Hatta, and Y. Koumoto for their technical assistance.

## Author contribution

TK, RN, and SSS conceived of the study and participated in its design and coordination. SSS, KO, and IHN designed the pMpGE vectors, and SSS, MS, JT, and YM constructed these vectors. RN designed the pMpGWB337 derivative vectors, and SI constructed these vectors. RN designed and RN, YM, and SI conducted the conditional knockout experiments. SSS and TS conducted the large deletion assays. YM conducted most of the other experiments. TK, RN, and SSS wrote the manuscript with inputs from co-authors.

---

**Figure S1.**
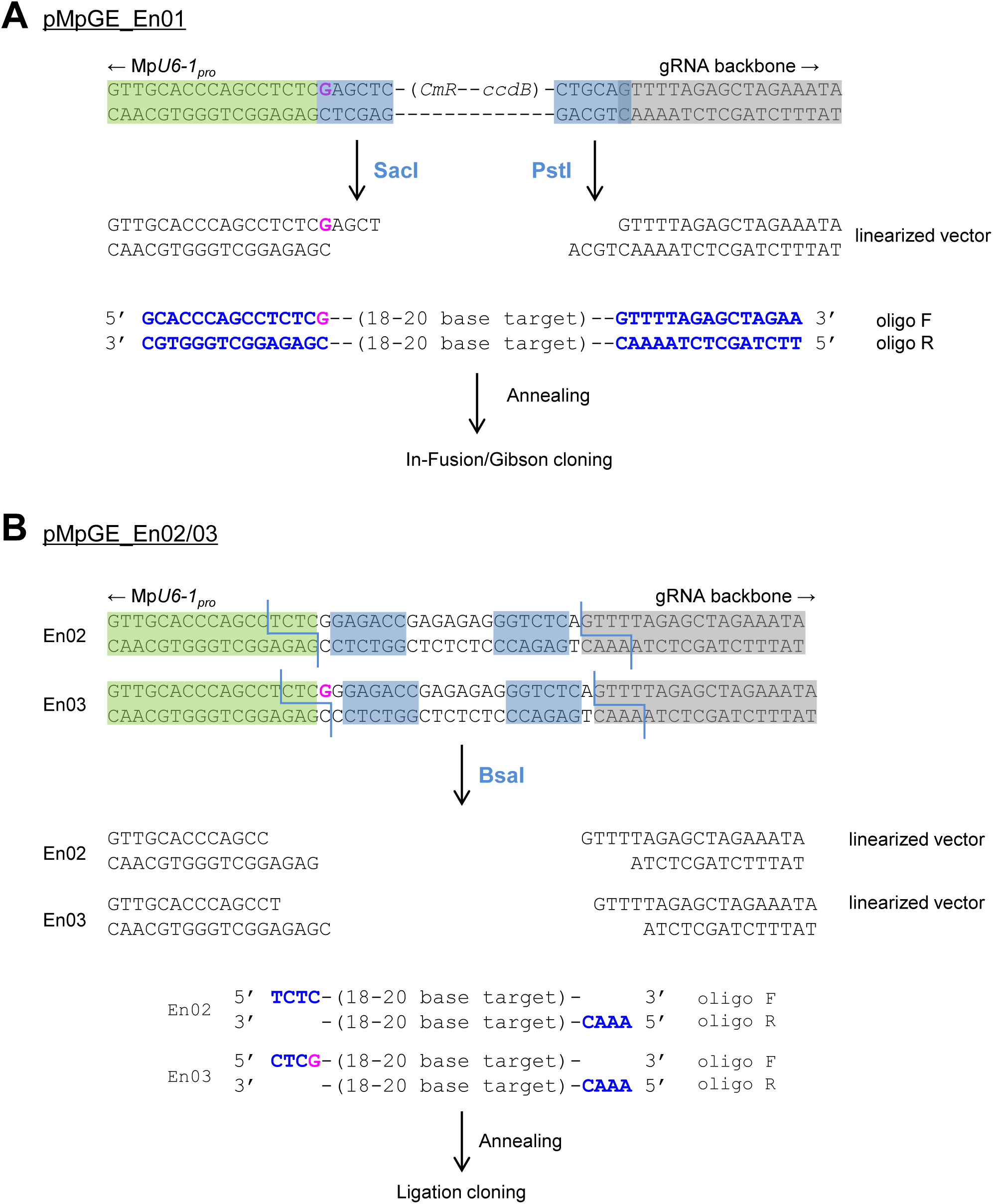
Protocols for gRNA cloning in entry vectors. (A) pMpGE_En01 is designed to use the In-Fusion/Gibson cloning methods. pMpGE_En01 is digested by SacI and PstI. Two entirely complementary oligo DNAs, which contain at both ends the 15-bp sequences identical to each end of the digested vector and a guide sequence without PAM sequence in between, are annealed and cloned by use the In-Fusion/Gibson reaction. The sense strand of gRNAs should be coded in oligo F. The ‘extra initial G’ is colored in magenta. (B) pMpGE_En02/03 are designed to use ligation reactions. pMpGE_En02 or pMpGE_En03 is digested by BsaI, which digests outside of its recognition sites. Oligo F, which contains a sense-strand guide sequence with TCTC at its 5’ end, and oligo R, which contains the reverse-complement guide sequence with AAAC at its 5’ end, are annealed and cloned by ligation reaction to pMpGE_En02. pMpGE_En02 does not contain an ‘extra initial G’ and thus requires G or A in the first nucleotide position of a guide sequence for efficient expression. In case of pMpGE_En03, the ‘extra initial G’ exists in the vector. Therefore, oligo F should contain a sense-strand guide sequence with CTCG at its 5’ end for the construction of pMpGE_En03.

**Figure S2.**
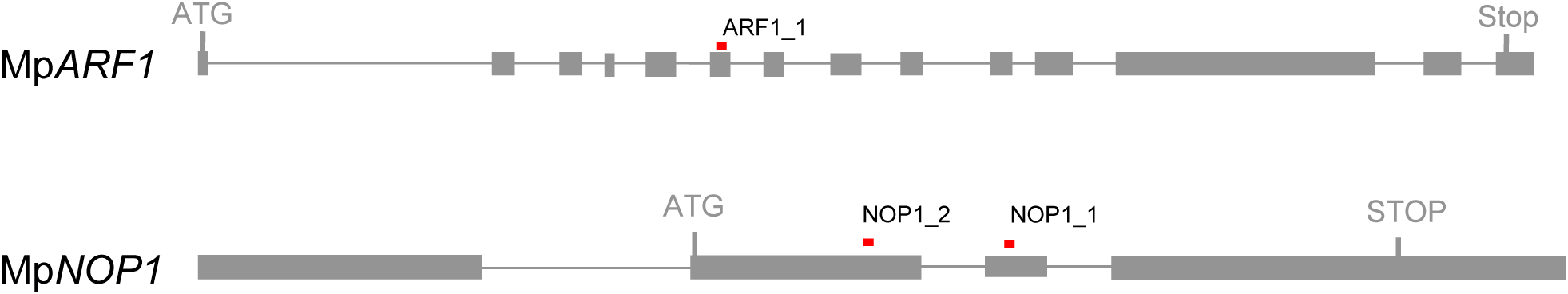
Gene structures of Mp*ARF1* and Mp*NOP1*. Target sites of the gRNAs used are shown as red lines. Boxes and lines show exons and introns, respectively. “ATG” and “Stop” denote the predicated initiation and termination codons.

**Figure S3.**
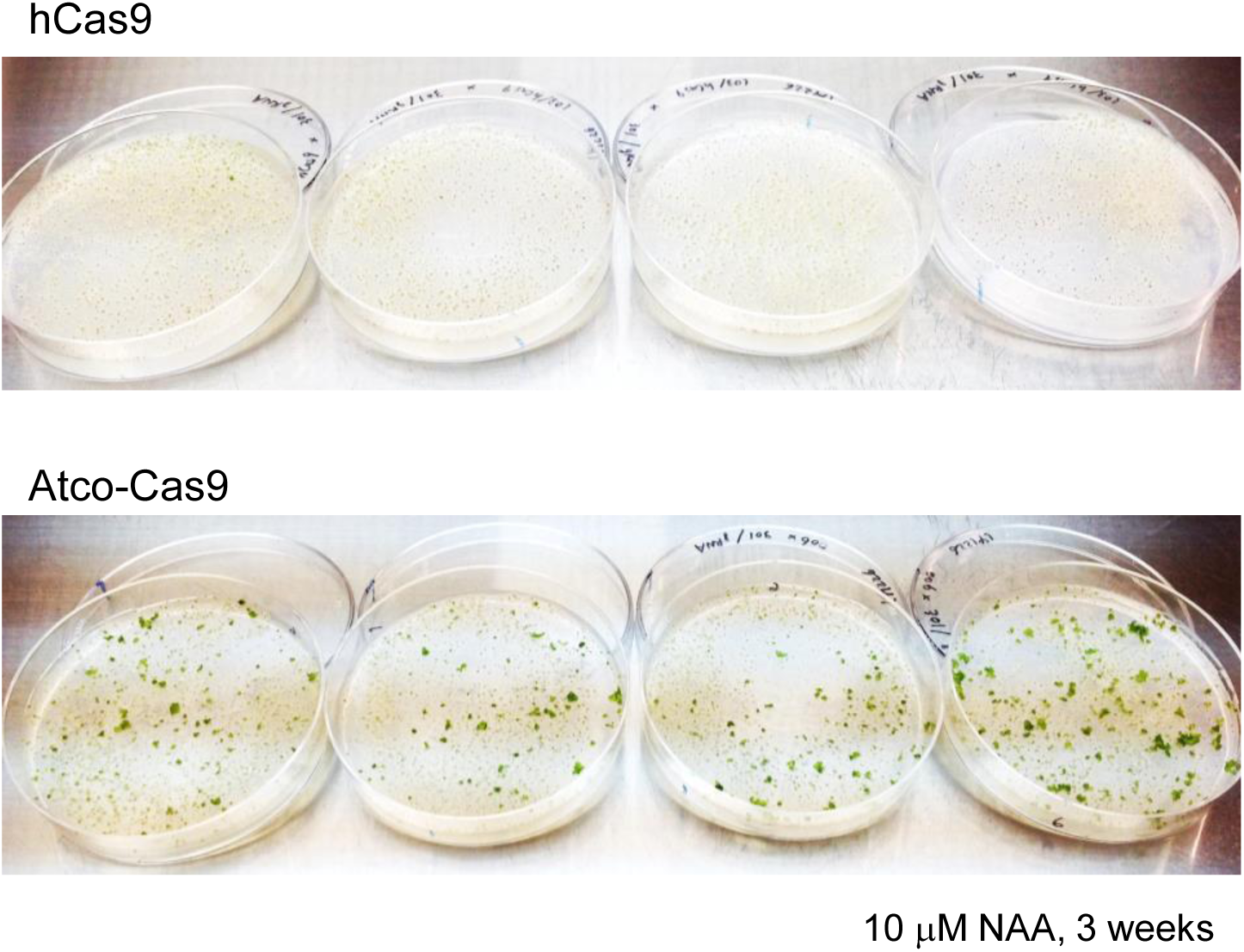
Comparison between *hCas9* and *Atco-Cas9* for genome editing efficiency. The same Mp*ARF1*-trageting gRNA expression vector (pMpGWB301_ARF1_1) was introduced into sporelings together with either hCas9 expression vector (pMpGWB103-hCas9; top) or Atco-Cas9 expression vector (pMpGE006; bottom) and selected on media containing 10 µM NAA for three weeks. Mutations in the Mp*ARF1* gene are known to cause NAA resistance.

**Figure S4.**
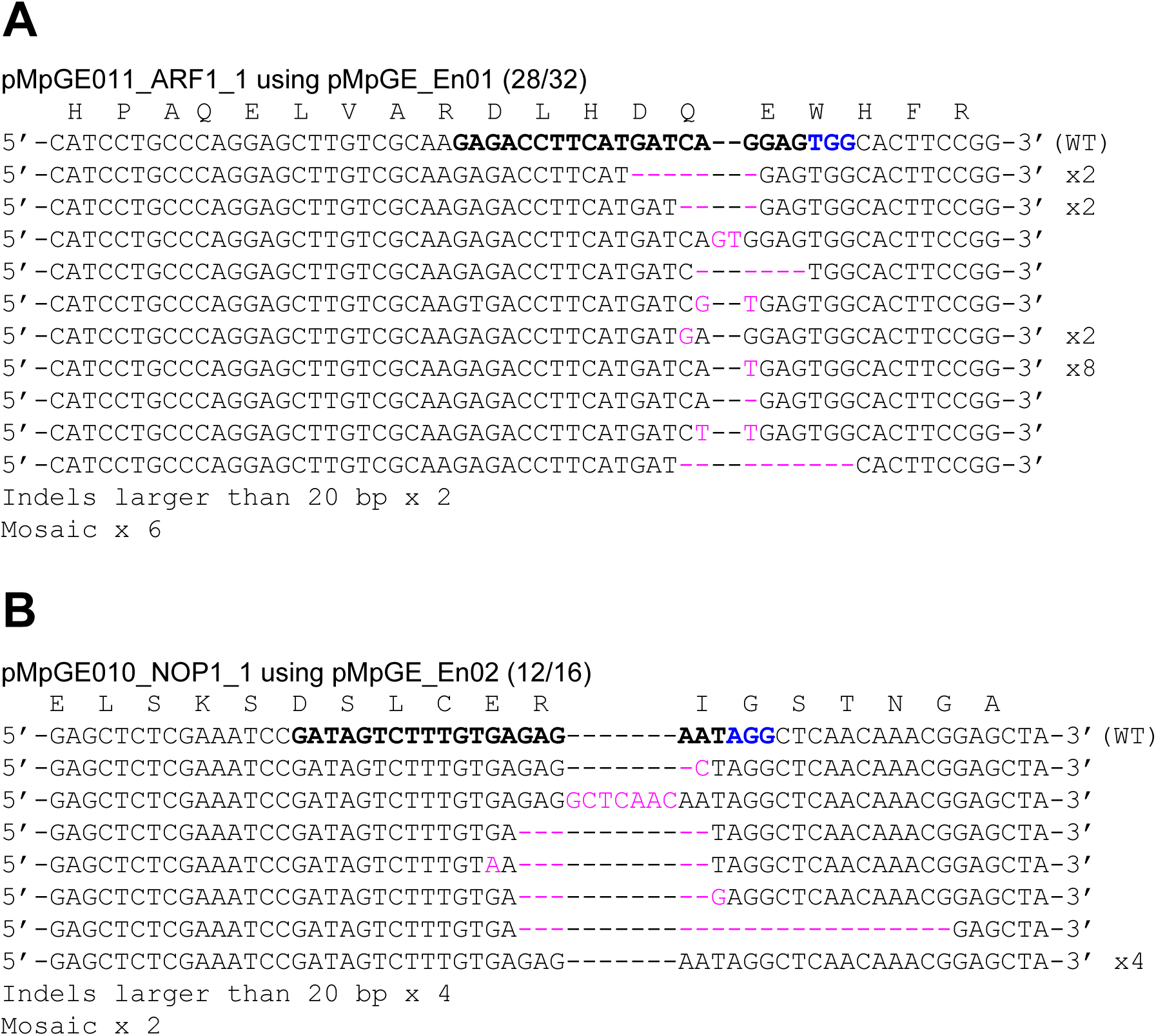
Genome editing with different vector combinations. (A)Mp*ARF1-*targeted mutagenesis with pMpGE011 containing the *gRNA* expression cassette for ARF1_1 derived from pMpGE_En01. (B)Mp*NOP1*-targeted mutagenesis with MpGE010 containing the *gRNA* expression cassette for NOP1_1 derived from pMpGE_En02. Inserted or substituted bases are colored in magenta. The target guide sequences are shown in bold face with their PAM sequences in blue.

**Figure S5.**
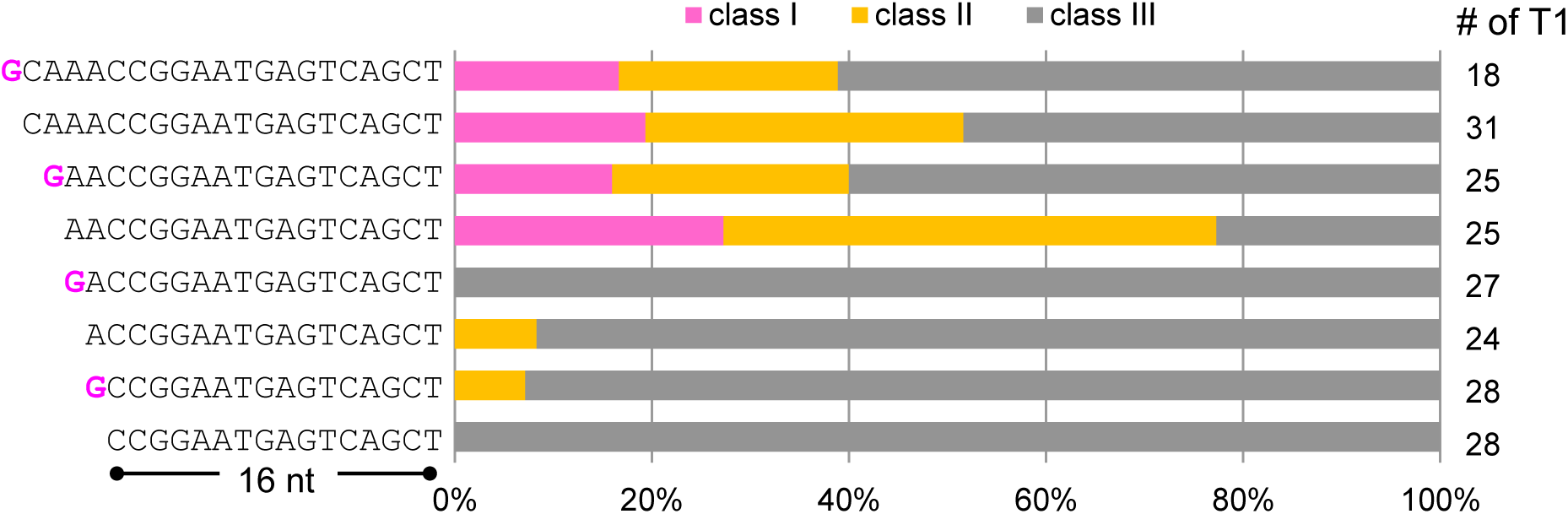
Effects of addition of an ‘extra initial G’. Proportions of mutant phenotype classes (see Fig. 4) in T1 plants transformed with Mp*NOP1*-targeting gRNAs (NOP1_2) of the indicated lengths with or without the ‘extra initial G’ (magenta). The numbers of T1 plants inspected are shown on the right side of the graph.

**Figure S6.**
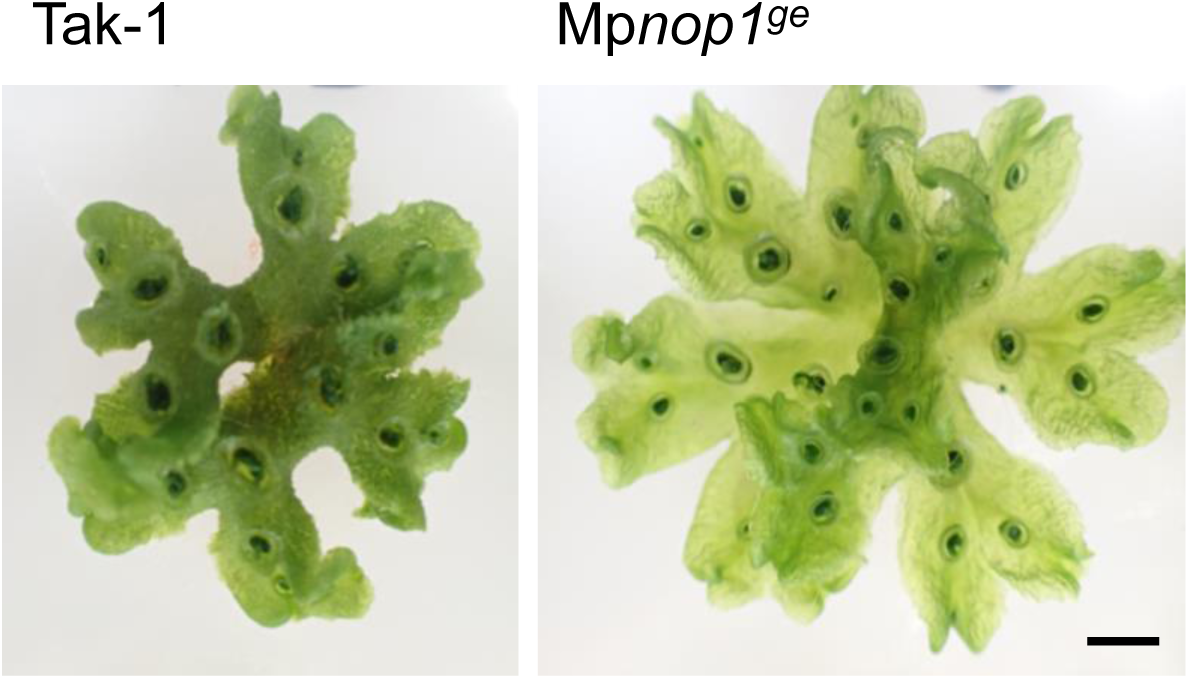
Representative photo of Mp*nop1*^ge^ mutant which harbors 4.5 kbp large deletion. Photos of Tak-1 control plant (Left) and the double transformant, which pMpGE013 with NOP1_3 gRNA and pMpGE014 with NOP1_6 gRNA were transfected (Right). Scale bar = 2 mm.

**Figure S7.**
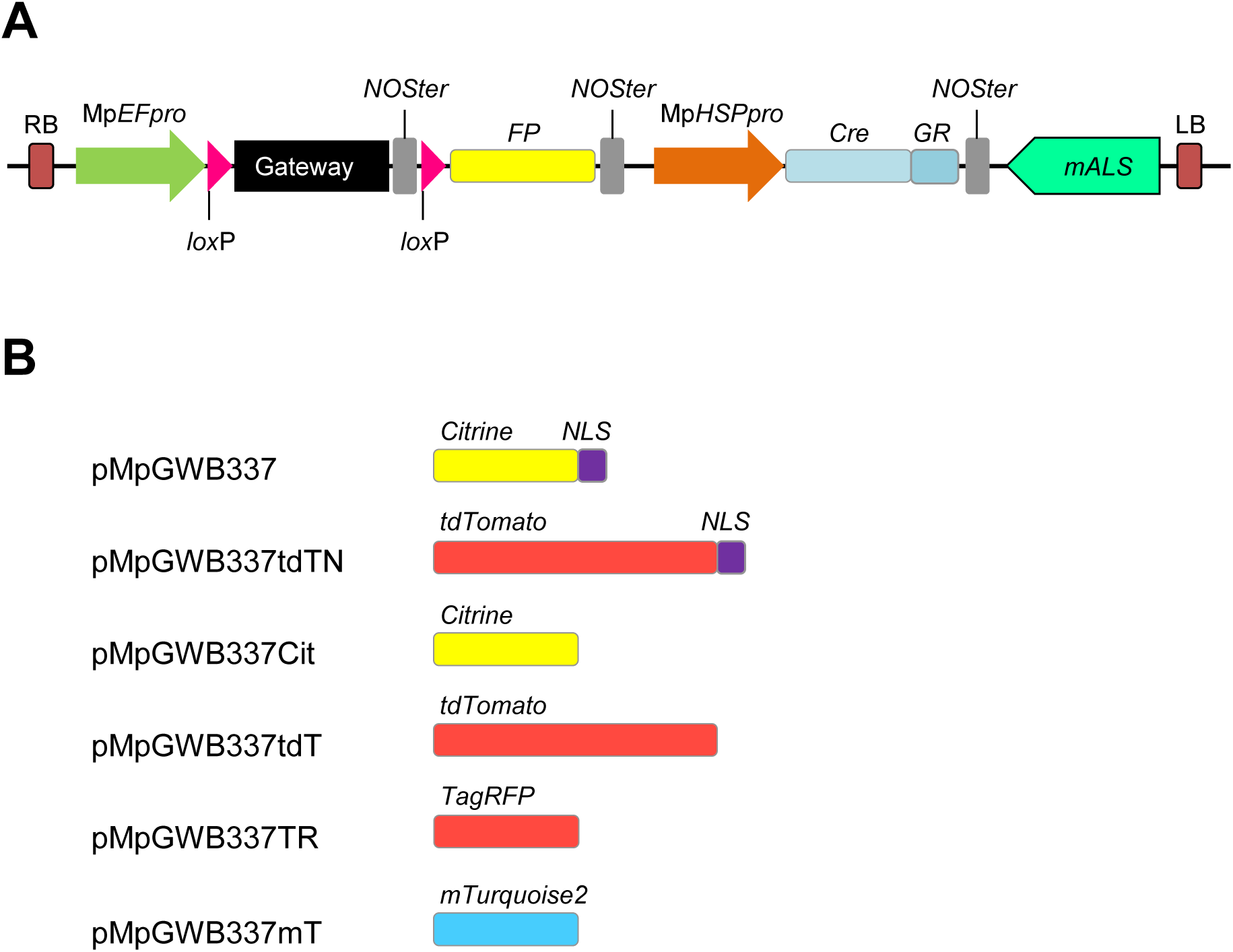
Derivatives of pMpGWB337 with various fluorescent protein markers. (A) Structure of pMpGWB337 derivatives. Genes for complementation (either cDNA or genomic fragment) can be expressed under the control of Mp*EF*_*pro*_ by introduction into the Gateway cassette and deleted in plants by heat shock and DEX treatment by virtue of the cassette expressing Cre recombinase fused to the rat glucocorticoid receptor domain (GR) under the control of the Mp*HSP17.8A1* promoter [23]. FP, fluorescent protein coding sequence. (B) List of fluorescent protein sequences in pMpGWB337 derivatives. NLS, nuclear localization signal.

**Figure S8.**
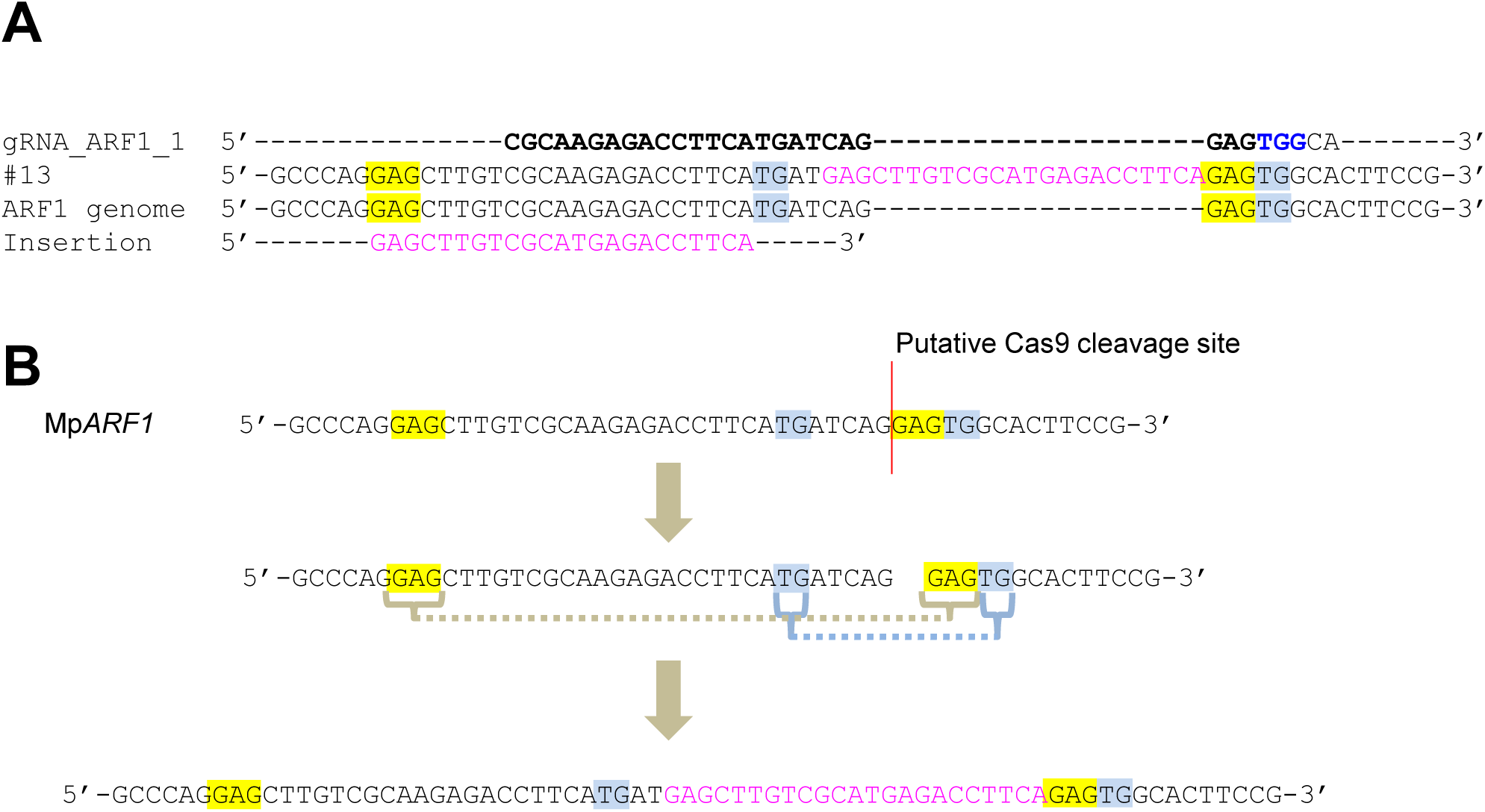
One of the example of microhomology-based repair in *M. polymorpha*. (A) Alignment of gDNA sequence of #13. Microhomology was highlighted with yellow and blue. (B) Schematic putative repair pathway in the mutant # 13 in pMpGE011_ARF1_1. The sequence was the same as Fig. S4

**Supplementary Table S1:**
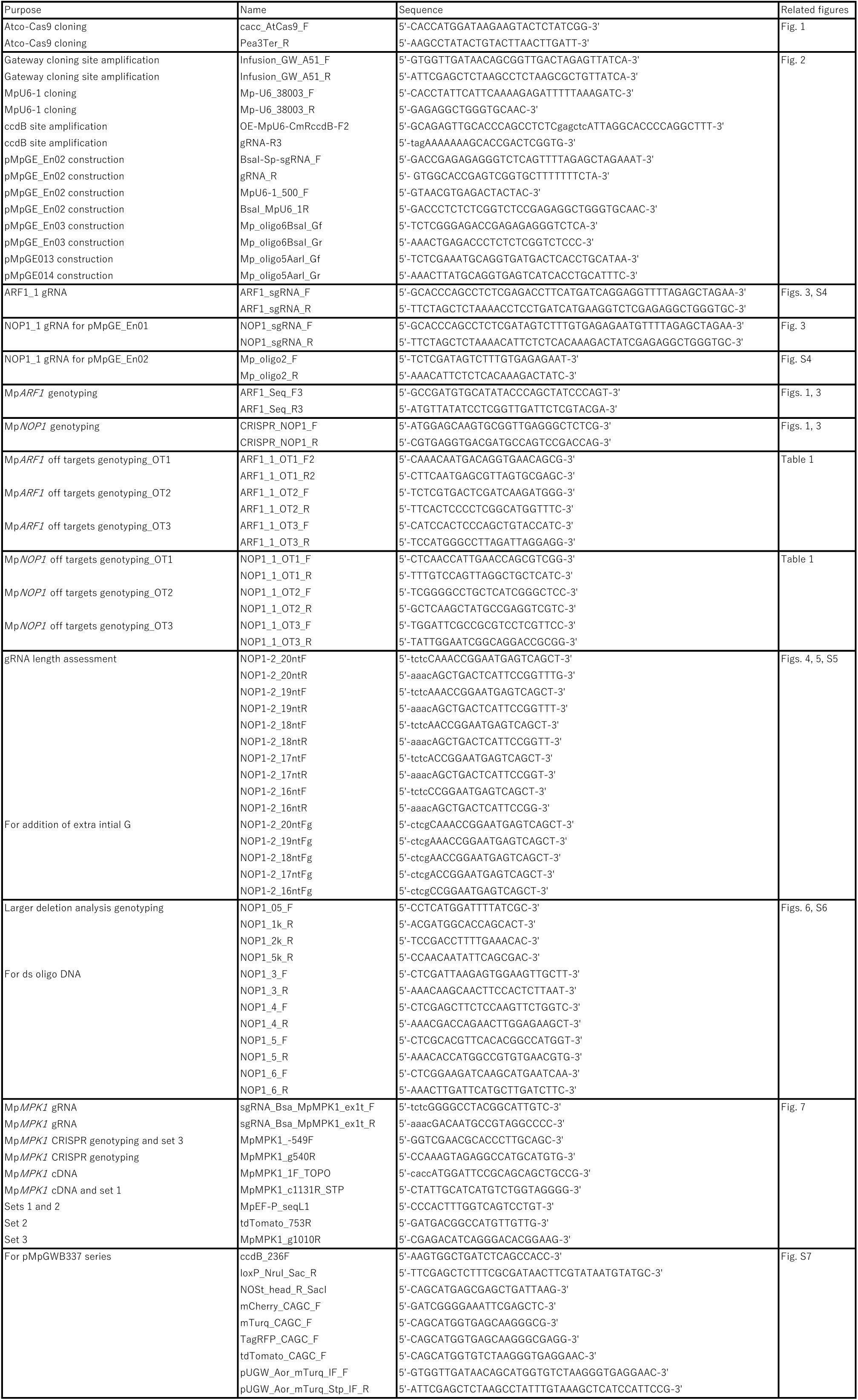
**Oligos used in this study**

## Abbreviations

ARF1: *AUXIN RESPONSE FACTOR1*
Cas9: CRISPR-associated endonuclease 9
CRISPR: clustered regularly interspaced short palindromic repeats
DSB: double-strand break
EF: *ELONGATION FACTOR1α*
HPT: *hygromycin phosphotransferase*
gRNA: single guide RNA
NAA: 1-naphthalene acetic acid
NHEJ: non-homologous end joining
NLS: nuclear localization signal
NOP1: *NOPPERABO1*
mALS: *mutated acetolactate synthase*
MMEJ: microhomology-mediated end joining
PAM: protospacer adjacent motif
PCR: polymerase chain reaction
RT-PCR: reverse transcription polymerase chain reaction.

